# Imaging transcriptomics: Convergent cellular, transcriptomic, and molecular neuroimaging signatures in the healthy adult human brain

**DOI:** 10.1101/2021.06.18.448872

**Authors:** Daniel Martins, Alessio Giacomel, Steven CR Williams, Federico Turkheimer, Ottavia Dipasquale, Mattia Veronese, PET templates working group

## Abstract

The expansion of neuroimaging techniques over the last decades has opened a wide range of new possibilities to characterize brain dysfunction in several neurological and psychiatric disorders. However, the lack of specificity of most of these techniques, such as magnetic resonance imaging (MRI)-derived measures, to the underlying molecular and cellular properties of the brain tissue poses limitations to the amount of information one can extract to inform precise models of brain disease. The integration of transcriptomic and neuroimaging data, known as ‘imaging transcriptomics’, has recently emerged as an indirect way forward to test and/or generate hypotheses about potential cellular and transcriptomic pathways that might underly specific changes in neuroimaging MRI biomarkers. However, the validity of this approach is yet to be examined in-depth. Here, we sought to bridge this gap by performing imaging transcriptomic analyses of the regional distribution of well-known molecular markers, assessed by positron emission tomography (PET), in the healthy human brain. We focused on tracers spanning different elements of the biology of the brain, including neuroreceptors, synaptic proteins, metabolism, and glia. Using transcriptome-wide data from the Allen Brain Atlas, we applied partial least square regression to rank genes according to their level of spatial alignment with the regional distribution of these neuroimaging markers in the brain. Then, we performed gene set enrichment analyses to explore the enrichment for specific biological and cell-type pathways among the genes most strongly associated with each neuroimaging marker. Overall, our findings show that imaging transcriptomics can recover plausible transcriptomic and cellular correlates of the regional distribution of benchmark molecular imaging markers, independently of the type of parcellation used to map gene expression and neuroimaging data. Our data support the plausibility and robustness of imaging transcriptomics as an indirect approach for bridging gene expression, cells and macroscopical neuroimaging and improving our understanding of the biological pathways underlying regional variability in neuroimaging features.

## Introduction

Over the past two decades, in-vivo human neuroimaging techniques, such as magnetic resonance imaging (MRI), have emerged as powerful tools to advance our understanding of macroscopic neural phenotypes measured across the entire brain^1^. The increasing application of MRI for studying neurological and psychiatric disorders has provided detailed anatomical characterizations of regional patterns of brain’s structural and functional alterations in these disorders^2,3^. However, the lack of specificity of most MRI-based techniques to the underlying molecular and cellular properties of the brain tissue^4^ has limited the potential of these neuroimaging biomarkers to inform mechanistic models of brain disease, understand biological mechanisms behind regional vulnerability to pathological changes or identify biological pathways that might be amenable to pharmacological intervention.

The recent introduction of comprehensive, brain-wide gene expression atlases such as the Allen Human Brain Atlas (AHBA) has opened new opportunities for understanding how spatial variations on the molecular transcriptomic scale relate to the macroscopic neuroimaging phenotypes^5,6^. This unprecedented capacity to link molecular pathways to macroscale brain organization has given rise to the emergent field of imaging transcriptomics^7^. Imaging transcriptomics is concerned with the identification of spatial correlations between gene expression patterns and some property of brain structure or function, as measured by neuroimaging^7^. The main goal of this approach is to identify genes with spatial profiles of regional expression that track anatomical variations in a certain neuroimaging biomarker. Typically, these analyses include mapping both gene expression data from the AHBA and neuroimaging maps to a common neuroimaging space (e.g., parcellated atlas of the brain). Then, one or multiple neuroimaging biomarkers (response variables) are related to expression measures of several thousands of genes in each region (predictor variables), often by using multivariate statistical techniques such as partial least square regression (PLS). As a result, genes are ranked according to the degree of spatial alignment of their expression with the neuroimaging biomarker under exam. An enrichment analysis is then performed on the top-ranking genes: when a significant number of top-ranking genes have a particular gene annotation (e.g., biological or molecular pathway) relative to the number of annotations present in a reference set (e.g., the entire genome), then the top-ranking genes are said to be enriched for that annotation. Because the top-ranking genes are strongly associated with the brain map of interest, the enriched annotations are used as an indirect way to test and generate hypotheses about the potential cellular and biological pathways that might underly specific neuroimaging features^7^. This approach has already begun to provide insights into how regional variations in gene expression relate to diverse properties of brain structure^8-13^ and function^14-20^, changes during brain disease^21-31^ or development^32-34^.

As the field develops, it is important, on the one side, to establish methodological guidelines to ensure consistent and reproducible results; and, on the other side, to examine the validity of this approach to capture indirect associations between gene expression, cells and macroscopical neuroimaging features. Recent efforts focusing on the first aspect have provided key tools and practical guides for the implementation of these analyses^7,35-37^, while the second aspect on validity is yet to be assessed in-depth. To the best of our knowledge, only two previous works have examined it within a limited scope. In both studies, maps of the mean expression of sets of genes related to oligodendrocytes were found to be positively correlated with MRI measures sensitive to myelin (i.e magnetization transfer ratio or T1W/T2W ratio)^8,31^. Whether imaging transcriptomics can also successfully recover plausible transcriptomic and cellular correlates of the regional distribution of neuroimaging biomarkers beyond those related to myelin is currently elusive.

In this work, we sought to bridge this gap by performing imaging transcriptomic analyses of the regional distribution of well-known molecular markers (as assessed by positron emission tomography, PET) in the healthy human brain. Thanks to a large collaborative effort, we could examine a vast number of tracers spanning different elements of the biology of the brain, including neuroreceptors, synaptic proteins, metabolism, and glia. While most PET tracers do not necessarily have binding affinity for a single specific brain cell-type or biological pathway, departing from a molecular neuroimaging phenotype where the target is known allows for a more precise generation of hypotheses regarding pathways that should be captured by imaging transcriptomics if the approach is valid (see Table 1 for a summary of all markers used and our respective *a priori* specific hypotheses regarding biological and cell-type pathways enrichment).

**Table 1.** List of neuroimaging markers and respective hypotheses. In this table, we present a summary of all neuroimaging markers used in our transcriptomics analyses and the respective *a priori* hypotheses we formulated in respect to the biological and cellular pathways that we expected to align with the regional distribution of each tracer.

| Domain | Marker | Main target | Hypothesis |
| --- | --- | --- | --- |
| Neuroreceptors, synaptic proteins and metabolism | [<sup>11</sup>C]Flumazenil | GABA<sub>A</sub> receptor | Alignment with the regional distribution of genes involved in synaptic structure and neurotransmission, which are highly expressed in populations of neuronal cells. |
|  | [<sup>18</sup>F]GE179 | NMDA receptor |  |
|  | [<sup>11</sup>C]UCB-J | Synaptic vesicle glycoprotein 2A (SV2A) |  |
|  | [<sup>18</sup>F]FDG | Fluorodeoxyglucose (metabolic activity) |  |
| Astroglia and myelin | [<sup>11</sup>C]BU99008 | Imidazoline<sub>2</sub> binding site (I<sub>2</sub>BS) | Alignment with the regional distribution of astrocytic genes. |
|  | L-[<sup>11</sup>C]deprenyl-D2 | Monoamine oxidase B (MAO-B) |  |
|  | Magnetization transfer ratio (MT) | Sensitive to myelin (including intra-cortical myelination) | Alignment with the regional distribution of oligodendrocytes and oligodendrocyte precursor cells. |
|  | Myelin Water Content (WC) | Sensitive to myelin (including intra-cortical myelination) |  |
| 18-kDa Translocator protein (TSPO) and cyclooxygenase (Cox) | [<sup>11</sup>C]PK11195 | TSPO<br>(first-generation) | Alignment with the regional distribution of genes of the neuroimmune response axis (microglia/astrocytes). |
|  | [<sup>18</sup>F]DPA174 | TSPO<br>(second-generation) |  |
|  | [<sup>11</sup>C]PBR28 |  |  |
|  | [<sup>11</sup>C]ER176 |  |  |
|  | [<sup>11</sup>C]PS13 | COX-1 |  |

## Methods

### Neuroimaging data

We capitalized on collaborations with several research groups (also referred as “PET templates working group”) to gather a large pool of templates of different PET tracers, including: [^11^C]Flumazenil^38^, [^18^F]GE179^39^, [^11^C]UCB-J (unpublished data), [^18^F]FDG (unpublished data), [^11^C]BU99008^40^, L-[^11^C]deprenyl-D2^41^, [^11^C]PK11195^42^, [^18^F]DPA174 (unpublished), [^11^C]PBR28^43^, [^11^C]ER176^44^ and [^11^C]PS13^45^. For each tracer, neuroimaging templates were created as average maps from independent sets of healthy controls (number of subjects ranging from 7 to 38). Data were acquired with different experimental design protocols and quantified in accordance to the best PET imaging practice for each of these radioligands, as described in the respective original studies. We also included two maps from MRI measures sensitive to myelin, namely a map of magnetization transfer ratio (MT) ^46^ and one of myelin water content (WC)^47^ for comparison with previous studies. For details on the samples, data acquisition and quantification and voxel-wise maps of each tracer please see Supplementary.

For each neuroimaging marker, we calculated the mean distribution within each of the 83 regions-of-interest of the Desikan-Killiany^**48**^ (DK) atlas by using the FMRIB software library (FSL, v6)^49^. Then, we calculated pairwise Pearson’s correlations between the normalised regional distribution of each pair of markers, assessing the significance of each pair-wise correlation by using tests that account for the inherent spatial autocorrelation of the data (see section below on spatial permutation testing) (Supplementary Figure S1).

In Figure 1 we provide a summary of our full imaging transcriptomics analysis pipeline (Figure 1), which we describe in detail below.

**Figure 1.**
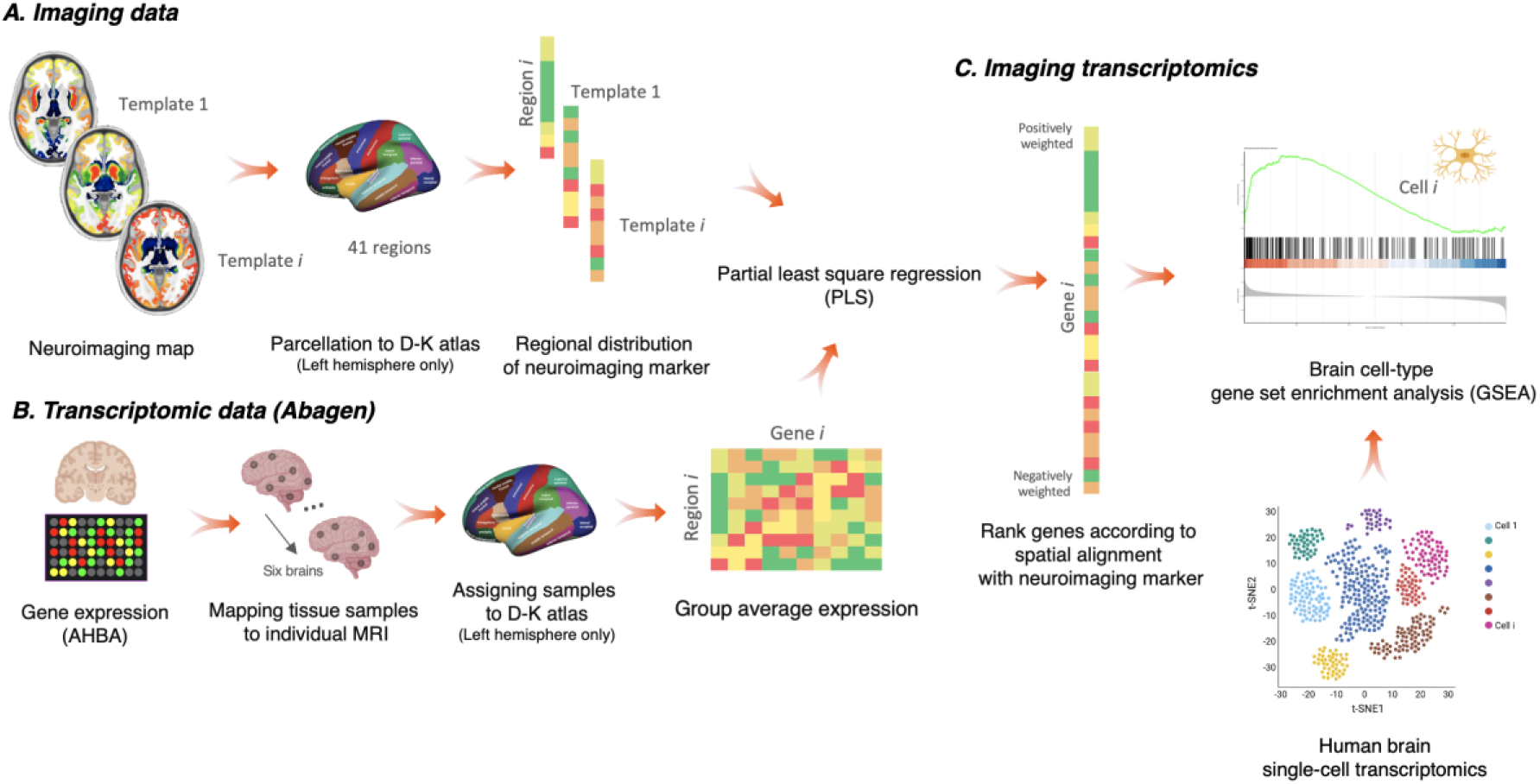
Imaging transcriptomics pipeline. (A) For each neuroimaging marker, we calculated the average distribution within each of the 83 regions-of-interest of the Desikan-Killiany (DK) atlas; only the data from the left hemisphere was used for further analyses since the Allen Human Brain Atlas only includes data from the right hemisphere for two subjects; (B) Gene expression analysis. We used *abagen* to obtain gene expression profiles from the AHBA in the 41 regions of the DK atlas (left hemisphere) across the six post-mortem brains sampled in this atlas. We excluded all genes with normalized expression values below the background (15,633 genes met this criterion). When more than one probe was available for a certain gene, we selected the probe with the highest consistency in expression across the 6 donors. We used partial least squares regression (PLS) to rank all genes according to their association with the regional distribution of each neuroimaging marker. Finally, we performed gene set enrichment analyses for gene ontologies and genes expressed in different cell-types (using single cell transcriptomic data from the human brain, as derived from previous studies). The gene set enrichment analyses were implemented with WebGestalt.

### Transcriptomic data

Regional microarray expression data were obtained from six post-mortem brains as part of the AHBA (http://human.brain-map.org/) (ages 24–57 years)^50^. We used the *abagen* toolbox (https://github.com/netneurolab/abagen) to process and map the transcriptomic data onto the 83 parcellated brain regions from the DK atlas. Briefly, genetic probes were reannotated using information provided by *Arnatkeviciute et al*., *2019*^7^ instead of the default probe information from the AHBA dataset, hence discarding probes that cannot be reliably matched to genes. Following previously published guidelines for probe-to-gene mappings and intensity-based filtering^7^, the reannotated probes were filtered based on their intensity relative to background noise level; probes with intensity lower than the background in ≥50% of samples were discarded. A single probe with the highest differential stability (highest pooled correlation across donors) was selected to represent each gene^51^. This procedure retained 15,633 probes, each representing a unique gene.

Next, tissue samples were assigned to brain regions using their corrected MNI coordinates (https://github.com/chrisfilo/alleninf) by finding the nearest region within a radius of 2 mm. To reduce the potential for misassignment, sample-to-region matching was constrained by hemisphere and cortical/subcortical divisions. If a brain region was not assigned to any sample based on the above procedure, the sample closest to the centroid of that region was selected to ensure that all brain regions were assigned a value. Samples assigned to the same brain region were averaged separately for each donor. Gene expression values were then normalized separately for each donor across regions using a robust sigmoid function and rescaled to the unit interval. We applied this procedure for cortical and subcortical regions separately, as suggested by *Arnatkeviciute et al*., *2019*^**7**^. Scaled expression profiles were finally averaged across donors, resulting in a single matrix with rows corresponding to brain regions and columns corresponding to the retained 15,633 genes. As a further robustness test, we conducted leave-one-donor out sensitivity analyses to generate six expression maps containing gene expression data from all donors, one at a time. The principal components of these six expression maps were highly correlated (average Pearson’s correlation of 0.993), supporting the idea that our final gene expression maps where we averaged gene expressions for each region across the six donors is unlikely to be biased by data from a specific donor. Since the AHBA only includes data for the right hemisphere for two subjects we only considered the 41 regions of left hemisphere regions (34 cortical plus 7 subcortical regions) for the following analyses.

### Partial least square regression

Partial least square regression uses the gene expression measurements (the predictor – or independent – variables) to predict regional variation in neuroimaging features (the response – or dependent – variables). This approach allows us to rank all genes by their multivariate spatial alignment with the regional distribution of each neuroimaging feature, while accounting for inherent collinearity in the predictor variables due to gene co-expression. The first PLS component (PLS1) is the linear combination of the weighted gene expression scores that have a brain expression map that covaries the most with the neuroimaging map. As the components are calculated to explain the maximum covariance between dependent and independent variables, the first component does not necessarily need to explain the maximum variance in the dependent variable. Of note, as the number of PLS components calculated increases, the amount of variance explained by each of them progressively decreases. Here, we examined models across a range of components (between 1 and 15) and evaluated the relative variance explained by each component against 1,000 null models, where we preserved the spatial autocorrelation of the data (see the section ‘spatial permutation test’ for more details). We then moved to further analyses only the PLS component explaining the largest amount of variance, which in our case was always the first component (PLS1). The error in estimating each gene’s PLS1 weight was assessed by bootstrapping (resampling with replacement of the 41 brain regions), and the ratio of the weight of each gene to its bootstrap standard error was used to calculate the *Z* scores and, hence, rank the genes according to their contribution to the PLS component under investigation^52^. Genes with large positive PLS weights correspond to genes that have higher than average expression in regions where the neuroimaging biomarker has a higher distribution, and lower than average expression in regions where the biomarker has a lower distribution. Mid-rank PLS weights showed expression gradients that are weakly related to the pattern of regional distribution of the neuroimaging biomarker. On the other side, genes with large negative PLS weights correspond to genes that have higher than average expression in regions where neuroimaging biomarker is lowly distributed, and lower than average expression in regions where the biomarker is highly distributed. Since in our analyses we were mostly interested in genes that track the distribution of each neuroimaging biomarker (i.e., that correlate positively with the biomarker), we focused on genes with positive PLS weights (even if we also present the results for genes with negative PLS weights for the curious reader).

### Spatial permutation test (spin test)

We assessed the significance of correlation between the neuroimaging biomarker and the selected PLS component using spatial permutation testing (spin test) to account for the inherent spatial autocorrelation of the imaging data, as implemented in previous studies^53-55^. This approach consists in comparing the empirical correlation amongst two spatial maps to a set of null correlations, generated by randomly rotating the spherical projection of one of the two spatial maps before projecting it back onto the brain parcel. Importantly, the rotated projection preserves the spatial contiguity of the empirical maps, as well as the hemispheric symmetry. Past studies using the spin test have focused on comparisons between brain maps including only cortical regions. However, subcortical regions were also of interest in this study. Since subcortical regions cannot be projected onto the inflated spherical pial surface, we incorporated the subcortex into our null models by shuffling the seven subcortical regions with respect to one another, whereas the cortical regions were shuffled using the spin test.

### Gene set enrichment analyses

We used each neuroimaging biomarker-associated ordered list of genes, ranked by the respective weights according to the PLS component under investigation, to perform gene set enrichment analyses (GSEA) for biological pathways (gene ontology) and genes expressed in different brain cell types, as identified in previous single-cell transcriptomic studies (see section on single-cell transcriptomic data below). We implemented these analyses using the GSEA method of interest of the Web-based gene set analysis toolkit (*WebGestalt*)^56^. In contrast to other over-representation methods, GSEA does not require the definition of an arbitrary threshold to isolate the most highly associated genes with a certain phenotype. It calculates an enrichment ratio (ER) that represents the degree to which the genes in the set are over-represented at either the top (positive ER) or bottom (negative ER) of the list, based on a Kolmogorov-Smirnov-like statistic. Then, it estimates the statistical significance of the ER using permutations to produce a null distribution for the ER (we used 1,000 permutations). The significance is determined by comparison to the null distribution. Finally, it adjusts for multiple hypothesis testing when several gene sets are being analysed at one time. The enrichment scores for each set are normalized (NER) to account for the number of genes in each gene set and a false discovery rate correction is applied.

### Single-cell transcriptomic data

To define the transcriptomic profile of the brain cell types mentioned above, we relied on the single-cell transcriptomic data from the previous single-cell transcriptomic study of Lake et al. ^57^. In this study, the authors used snDrop-seq Unique Molecular Identifier counts for cells from the visual (BA17) and dorsal frontal cortex (BA 6/9/10) and identified 30 brain cell-types (astrocytes, endothelial cells, pericytes, microglia, oligodendrocytes, oligodendrocytes progenitor cells (OPCs), 13 subtypes of excitatory neurons and 11 subtypes of inhibitory neurons). We decided to consider these 30 cells, since this classification provides a good coverage of a wide range of different neuronal and non-neuronal cells.

### Sensitivity analyses

We assessed the robustness of our findings by performing a sensitivity analysis examining the impact of: i) brain parcellation; ii) transcriptomic profile of brain cell-types; and iii) inclusion of subcortical + cortical vs cortical regions only, when a whole brain receptor neuroimaging marker (e.g. [^18^F]Fallypride) has higher binding in subcortical than cortical regions. These analyses are described in detail in Supplementary.

### Data availability

While some of these templates are publicly available, others have been kindly shared by individual research groups. Access to these data might be granted upon to reasonable request by contacting directly principal investigators (see Appendix with all contributors and associated data).

### Code availability

The code for performing the imaging transcriptomic analyses is now available as a python package that can be downloaded from https://github.com/molecular-neuroimaging/Imaging_Transcriptomics.

## Results

We provide a summary of the pairwise correlations between all maps in Supplementary Figure S1. Of note, we found large correlations between the regional distribution of [^11^C]BU99008 and L-[^11^C]deprenyl-D2 (r=0.90, p_spin_ <0.001), and between [^11^C]PBR28 and [^11^C]ER176 (r=0.99, p_spin_ <0.001). Below, we present the results of the imaging transcriptomics analyses for each map, organized in three main subsections: i) neuroreceptors, synaptic proteins and metabolism; ii) astroglia and myelin; iii) 18-kDa Translocator protein (TSPO) and Cox-1.

### Neuroreceptors, synaptic proteins and metabolism

#### [^11^C]Flumazenil

The first PLS component explained the largest amount of variance (44.99%) and correlated positively with the regional distribution of [^11^C]Flumazenil (r=0.6708, p_spin_ < 0.001) (Figure 2). We found enrichment among the most positively weighted genes for several gene ontology – biological process domain terms globally related to synaptic structure and transmission, including the glutamate receptor signalling pathway (NER=2.26, p_FDR_ = 0.001) and GABAergic synaptic transmission (NER=1.809, p_FDR_ = 0.02) (Supplementary data S2). The cell-type enrichment analysis indicated significant enrichment for several subclusters of excitatory and inhibitory neurons, with Ex6b emerging as the strongest enrichment hit. In addition to these neuronal subclusters, we also found significant enrichment for genes expressed in astrocytes and OPCs.

**Figure 2.**
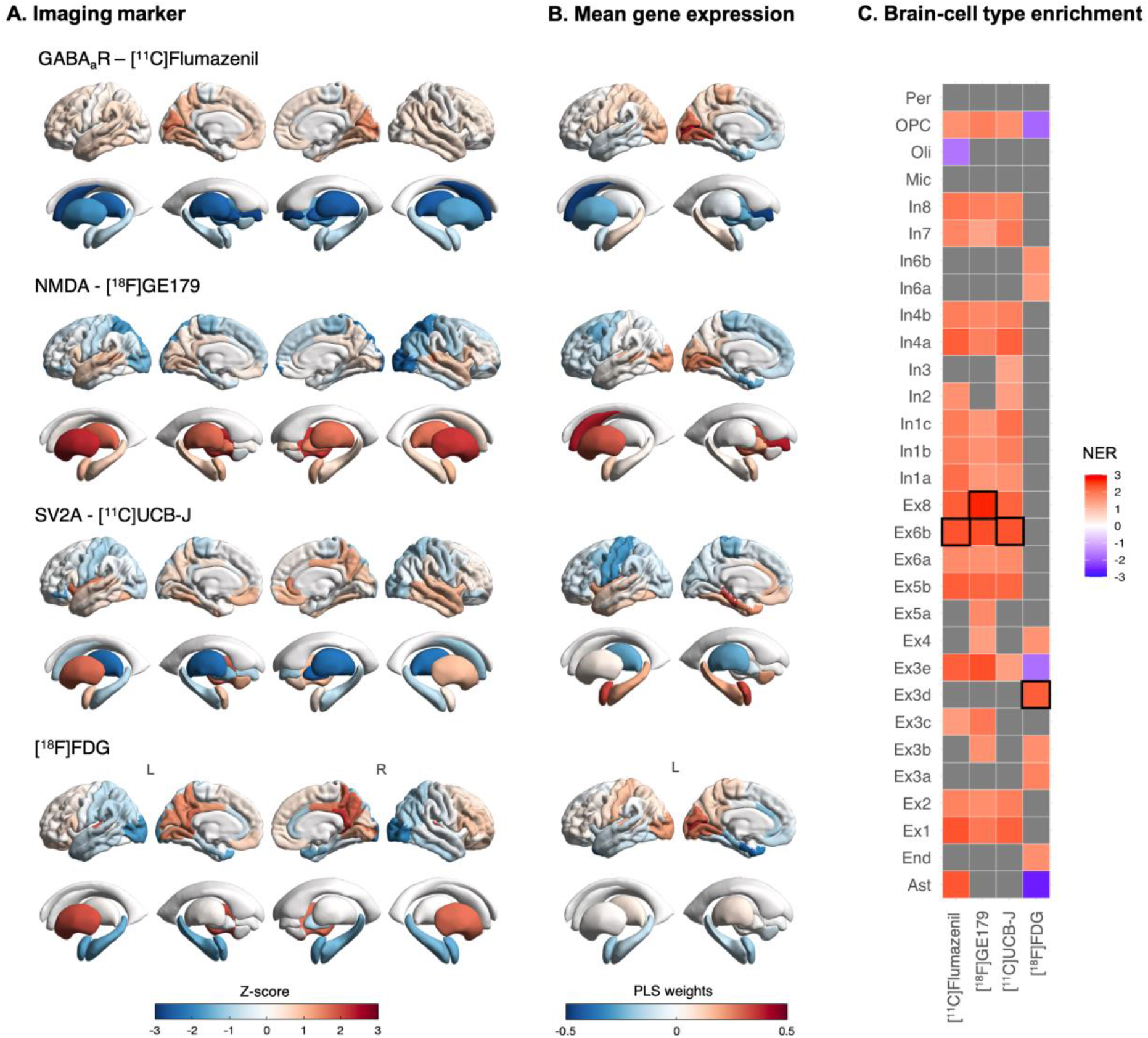
Imaging transcriptomics (neuroreceptors, synaptic proteins and metabolism). (A) - Regional distribution of each marker. (B) – Regional distribution of the PLS1 weights. (C) – Brain cell-type enrichment analyses: Positive normalized enrichment ratios (NER) indicate enrichment for genes of a certain brain cell-type among the genes with positive weights in PLS1 (i.e. positively correlated with the distribution of the neuroimaging marker); negative NERs indicate the reverse; grey squares indicate that the NER did not reach significance (pFDR>0.05). The full statistics underlying the tile plot can be found in Supplementary data S1. Abbreviations: Ast – astrocytes; End – endothelial; Ex – excitatory neurons; In - inhibitory neurons; Mic – microglia; Oli – oligodendrocytes; OPC – oligodendrocyte precursor cells; Per – pericytes.

#### [^18^F]GE179

The first PLS component explained the largest amount of variance (30.29%) and correlated positively with the regional distribution of [^18^F]GE179 (r=0.5504, p_spin_ = 0.002) (Figure 2). We found enrichment among the most positively weighted genes for two gene ontology – biological process terms, namely the synapse organization (NER=2.07, p_FDR_ = 0.02) and GABA signalling pathway (NER=2.02, p_FDR_ = 0.02) (Supplementary data S2). The cell-type enrichment analysis indicated significant enrichment for several subclusters of excitatory and inhibitory neurons, with Ex8 emerging as the strongest enrichment hit.

#### [^11^C]UCB-J

The first PLS component explained the largest amount of variance (30.92%) and correlated positively with the regional distribution of [^11^C]UCB-J (r=0.5561, pspin = 0.002) (Figure 2). We found enrichment among the most positively weighted genes for several gene ontology – biological process terms, including the neuropeptide signalling pathway (NER=2.11, p_FDR_ = 0.007), regulation of synaptic structure and activity (NER=1.98, p_FDR_ = 0.02) and glutamatergic synaptic transmission (NER=1.98, p_FDR_ = 0.02) (Supplementary data S2). The cell-type enrichment analysis indicated significant enrichment for several subclusters of excitatory and inhibitory neurons, with Ex6b emerging as the strongest enrichment hit.

#### [^18^F]FDG

The first PLS component explained the largest amount of variance (42.28%) and correlated positively with the regional distribution of [^18^F]FDG (r=0.6503, p_spin_ < 0.001) (Figure 2). We found enrichment among the most positively weighted genes for two gene ontology – biological process terms, which were peptide catabolic process (NER=1.97, p_FDR_ = 0.004) and protein dealkylation (NER=1.81, p_FDR_ = 0.007) (Supplementary data S2). The cell-type enrichment analysis indicated significant enrichment for several subclusters of excitatory and inhibitory neurons, with Ex3d emerging as the strongest enrichment hit. Among the non-neuronal cells, we also found enrichment for genes expressed in endothelial cells.

### Astroglia and myelin

#### [^11^C]BU99008]

The first PLS component explained the largest amount of variance (42.01%) and correlated positively with the regional distribution of [^11^C]BU99008 (r=0.6482, p_spin_ < 0.001) (Figure 3). We found enrichment among the most positively weighted genes for several gene ontology – biological process terms, globally related to protein targeting, RNA and ribonucleoproteins metabolism, and the GABA signalling pathway (Supplementary data S2). The cell-type enrichment analysis indicated significant enrichment for some subclusters of inhibitory neurons and one subcluster of excitatory neurons. Astrocytes emerged as the strongest enrichment hit. In addition, we also found significant enrichment for genes expressed in OPCs and Pericytes.

**Figure 3.**
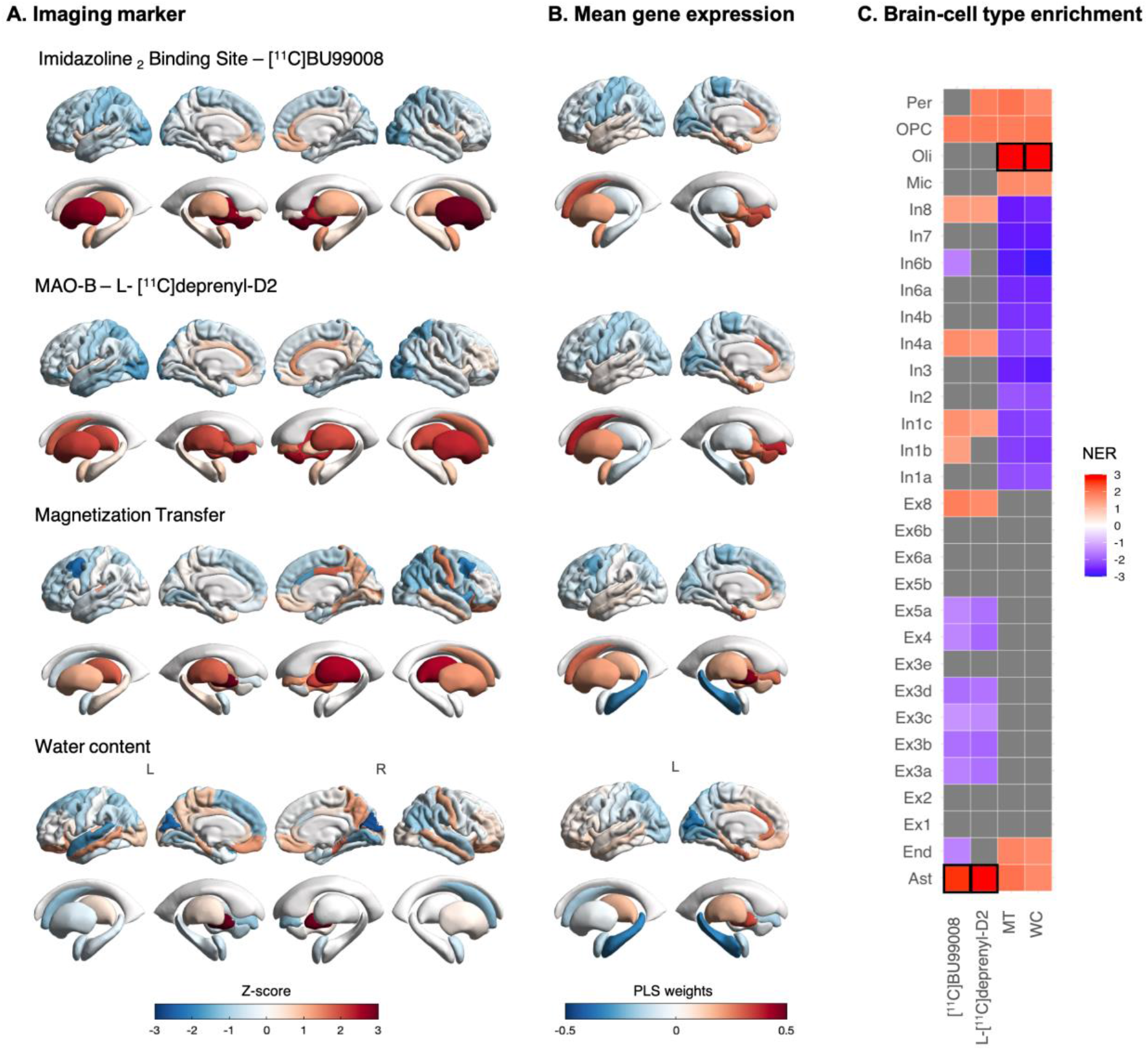
Imaging transcriptomics (astroglia and myelin). (A) - Regional distribution of each marker. (B) – Regional distribution of the PLS1 weights. (C) – Brain cell-type enrichment analyses: Positive normalized enrichment ratios (NER) indicate enrichment for genes of a certain brain cell-type among the genes with positive weights in PLS1 (i.e., positively correlated with the distribution of the neuroimaging marker); negative NERs indicate the reverse; grey squares indicate that the NER did not reach significance (pFDR>0.05); The full statistics underlying the tile plot can be found in Supplementary data S1. Abbreviations: Ast – astrocytes; End – endothelial; Ex – excitatory neurons; In - inhibitory neurons; Mic – microglia; Oli – oligodendrocytes; OPC – oligodendrocyte precursor cells; Per – pericytes.

#### L-[^11^C]deprenyl-D2

The first PLS component explained the largest amount of variance (41.47%) and correlated positively with the regional distribution of L-[^11^C]deprenyl-D2 (r=0.6440, p_spin_ < 0.001) (Figure 3). We found enrichment among the most positively weighted genes for several gene ontology – biological process terms, globally related to protein targeting, RNA and ribonucleoproteins metabolism (Supplementary data S2). The cell-type enrichment analysis indicated significant enrichment for some subclusters of inhibitory neurons and one subcluster of excitatory neurons. Astrocytes emerged as the strongest enrichment hit. In addition, we also found significant enrichment for genes expressed in OPCs.

#### Magnetization transfer ratio (MT)

The first PLS component explained the largest amount of variance (34.91%) and correlated positively with the regional distribution of MT (r=0.5909, p_spin_ < 0.001) (Figure 3). We found enrichment among the most positively weighted genes for several gene ontology – biological process terms, globally related to extracellular structure organization, endothelium development and immune response (Supplementary data S2). The cell-type enrichment analysis indicated significant enrichment for all non-neuronal cells. Oligodendrocytes emerged as the strongest enrichment hit.

#### Myelin water content (WC)

The first PLS component explained the largest amount of variance (31.40%) and correlated positively with the regional distribution of MT (r=0.5604, p_spin_ = 0.007) (Figure 3). We found enrichment among the most positively weighted genes for several gene ontology – biological process terms, globally related to amino acid biosynthetic processes, extracellular structure organization, endothelium development and angiogenesis, cell adhesion mediated by integrins and immune response (Supplementary data S2). The cell-type enrichment analysis indicated significant enrichment for all non-neuronal cells. Oligodendrocytes emerged as the strongest enrichment hit.

### 18-kDa translocator protein (TSPO) and cyclooxygenase (Cox)

#### TSPO - [^11^C]PK11195

The first PLS component explained the largest amount of variance (36.30%) and correlated positively with the regional distribution of [^11^C]PK11195 (r=0.6025, p_spin_ < 0.001) (Figure 4). We found enrichment among the most positively weighted genes for several gene ontology – biological process terms, including metabolic processes, granulocyte activation and response to interferon gamma, ensheathment of neurons, and DNA repair (Supplementary data S2). The cell-type enrichment analysis indicated significant enrichment for all non-neuronal cells, excluding astrocytes. Microglia emerged as the strongest enrichment hit.

**Figure 4.**
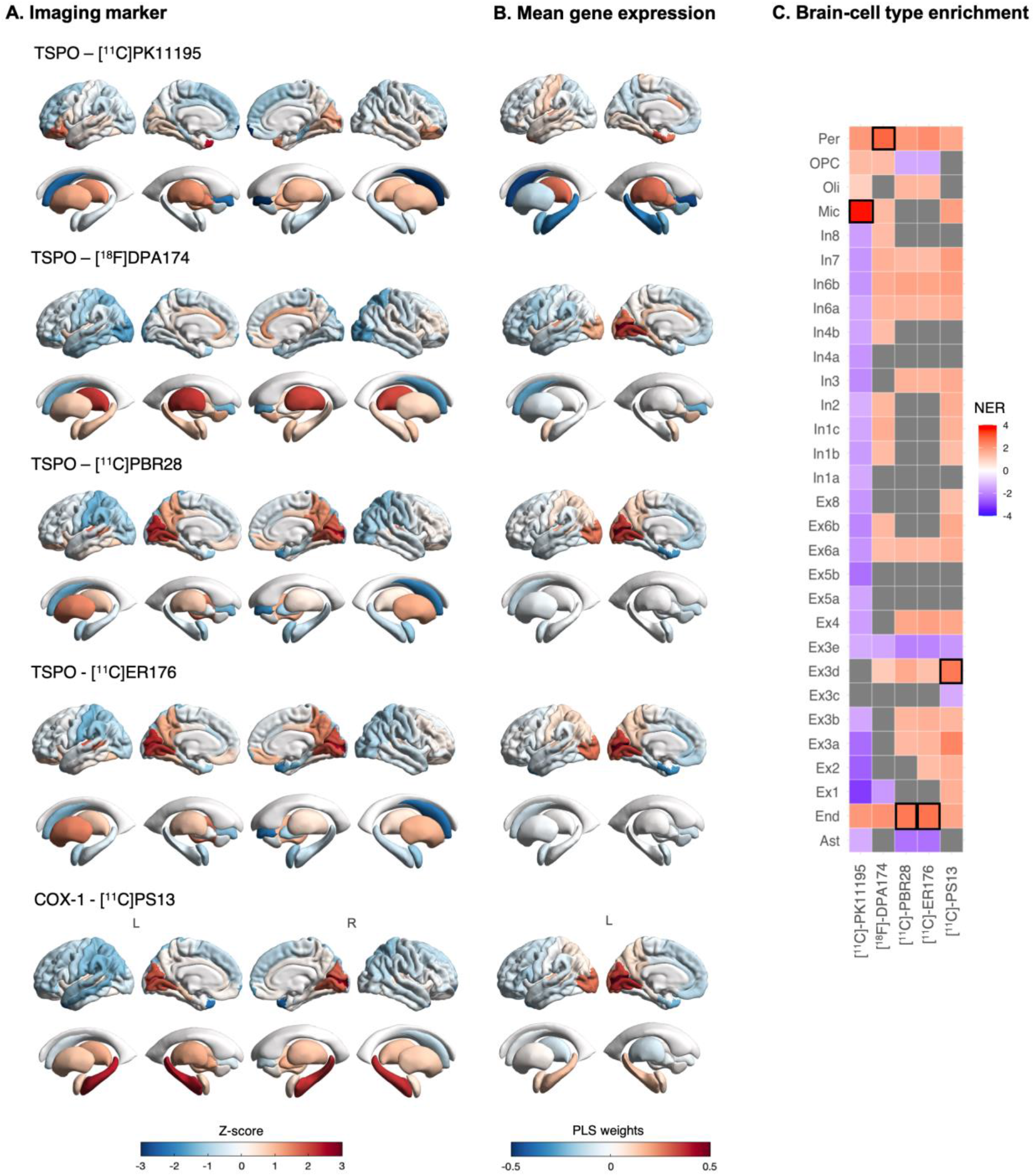
Imaging transcriptomics (Translocator protein and cyclooxygenase). (A) - Regional distribution of each marker. (B) – Regional distribution of the PLS1 weights. (C) – Brain cell-type enrichment analyses: Positive normalized enrichment ratios (NER) indicate enrichment for genes of a certain brain cell-type among the genes with positive weights in PLS1 (i.e. positively correlated with the distribution of the neuroimaging marker); negative NERs indicate the reverse; grey squares indicate that the NER did not reach significance (pFDR>0.05); The full statistics underlying the tile plot can be found in Supplementary data S1. Abbreviations: Ast – astrocytes; End – endothelial; Ex – excitatory neurons; In - inhibitory neurons; Mic – microglia; Oli – oligodendrocytes; OPC – oligodendrocyte precursor cells; Per – pericytes.

#### TSPO - [^18^F]DPA174

The first PLS component explained the largest amount of variance (33.23%) and correlated positively with the regional distribution of [^18^F]DPA174 (r=0.5765, p_spin_ = 0.001) (Figure 4). We did not find enrichment among the most positively weighted genes for any of the gene ontology – biological process terms. The cell-type enrichment analysis indicated significant enrichment for all non-neuronal cells, excluding astrocytes and oligodendrocytes. In contrast with [^11^C]PK11195, here we also found enrichment for several subclusters of excitatory and inhibitory cells. Pericytes emerged as the strongest enrichment hit.

#### TSPO - [^18^C]PBR28

The first PLS component explained the largest amount of variance (34.66%) and correlated positively with the regional distribution of [^18^C]PBR28 (r=0.5887, p_spin_ = 0.001) (Figure 4). We did not find enrichment among the most positively weighted genes for any of the gene ontology – biological process terms. The cell-type enrichment analysis indicated significant enrichment for pericytes, oligodendrocytes, endothelial cells and several subclusters of excitatory and inhibitory cells. Endothelial cells emerged as the strongest enrichment hit.

#### TSPO - [^18^C]ER176

The first PLS component explained the largest amount of variance (36.60%) and correlated positively with the regional distribution of [^18^C]ER176 (r=0.6050, p_spin_ < 0.001) (Figure 4). We did not find enrichment among the most positively weighted genes for any of the gene ontology – biological process terms. The cell-type enrichment analysis indicated significant enrichment for pericytes, oligodendrocytes, endothelial cells and several subclusters of excitatory and inhibitory cells. Endothelial cells emerged as the strongest enrichment hit.

#### COX-1 - [^11^C]PS13

The first PLS component explained the largest amount of variance (41.22%) and correlated positively with the regional distribution of [^11^C]PS13 (r=0.6420, p_spin_ < 0.001) (Figure 4). We did not find enrichment among the most positively weighted genes for any of the gene ontology – biological process terms. The cell-type enrichment analysis indicated significant enrichment for pericytes, microglia, endothelial cells and several subclusters of excitatory and inhibitory cells, with Ex3d emerging as the strongest enrichment hit

### Sensitivity analyses

#### Brain parcellation

We found large correlations (Pearson’s r between 0.79 and 0.94) between PLS1 weights of all genes as estimated using the DK and AAL3 atlases (Supplementary Table S1).

#### Brain cell-types transcriptomic profile

Changing the gene sets used to define the transcriptomic profile of each brain cell-type did not considerably change our main brain cell-type enrichment conclusions (Supplementary data S3).

#### Inclusion of cortical + subcortical regions vs cortical regions only

By including both cortical and subcortical regions in the same model, we identified the DRD2 gene as one of the top genes positively associated with [^18^F]fallypride binding (Z = 5.17, Rank: 161/15,633, top 1% of all genes). When repeating the same analysis with cortical regions, we still identified *DRD2* as a positively weighted gene, but its rank was considerably lower (Z = 3.67, Rank: 2,204/15,633, top 15% of all genes) (Supplementary Figure S2).

## Discussion

In this work, we investigated whether imaging transcriptomics can recover plausible transcriptomic and cellular correlates of the regional distribution of benchmark molecular imaging markers. With the recent upsurge of imaging transcriptomics studies, this work is crucial to strengthening our confidence in the plausibility of this integrative approach as an indirect way to generate hypotheses about potential biological pathways explaining regional variations in *in vivo* neuroimaging biomarkers. This is of particular importance for those neuroimaging markers where more fine-grained biological mechanisms cannot be measured directly. Thanks to an unprecedented collaborative effort, we gathered a vast number of tracers spanning different elements of the brain’s biology, which allowed us to conduct a comprehensive evaluation of the plausibility of the approach across different biological and cellular pathways. In a set of secondary analyses, we confirmed that the method is likely to generalize well beyond the specific choices of brain parcellation or gene sets used to define the transcriptomic profile of brain cell-types. Altogether, our data supports the plausibility and robustness of imaging transcriptomics as an indirect approach bridging levels between gene expression, cells and macroscopical neuroimaging to improve our understanding of the biological pathways underlying regional variability in neuroimaging features.

Our first set of analyses were focused on four markers targeting neuroreceptors (GABA-A and NMDA receptors), a synaptic protein (SV2A) and metabolism (FDG). We hypothesized that the regional distribution of these markers would align preferentially with the distribution of genes involved in synaptic structure and neurotransmission, which are highly expressed in populations of neuronal cells. In line with our main hypotheses, this was what we found. The GABA_A_ receptor is the major target for GABA, an inhibitory neurotransmitter found in about 20–50% of synapses in the brain ^58^. This receptor is predominantly located in the postsynaptic membrane and can be expressed by both excitatory and inhibitory neurons^59^; however, it also localizes at extra-synaptic sites. For instance, it has become increasingly clear that astrocytes, both in cell culture and in tissue slices, express abundant GABA_A_ receptors ^60^. This aspect might explain why, in addition to neuronal cells, we also found enrichment for astrocytic genes among the top genes positively associated with [^11^C]Flumazenil binding.

The NMDA receptor is a glutamate receptor predominantly located in the post-synaptic membrane of neurons, which participates in neuronal excitation and plasticity^61-63^. It is present at both excitatory and inhibitory neurons ^64,65^; but there are also reports of its expression in oligodendrocytes and OPCs^66,67^. While we found enrichment among the genes positively associated with NMDA regional distribution for several clusters of excitatory and inhibitory neurons and oligodendrocytes progenitors, we did not find enrichment for genes highly expressed in mature oligodendrocytes. This could reflect the fact that expression of NMDA receptors in oligodendrocytes is only minoritarian as compared to neurons, which might be insufficient to generate a pattern of distribution that would strongly align with oligodendrocyte genes.

The synaptic vesicle glycoprotein 2A (SV2A) is a prototype presynaptic vesicle protein regulating action potential-dependent neurotransmitters release, which is expressed across both excitatory and inhibitory neurons ^68,69^. This cellular distribution aligns well with the fact that we found enrichment among the genes positively associated with SV2A regional distribution for several clusters of excitatory and inhibitory neurons. To our knowledge, there have not been reports of high constitutive expression of SV2A in OPCs even if we found enrichment for genes highly expressed in these cells; however, since OPCs can also differentiate into neurons^70^, it is not implausible that SV2A might also be present in OPCs and colocalize with other elements of the molecular machinery of these cells.

[^18^F]FDG is a metabolic marker typically used as a fully quantitative indirect index of neuronal activity as it captures the cerebral glucose uptake^71,72^. A long-standing model postulates that the use of glucose by neurons requires an interplay with astrocytes in an astrocyte-to-neuron lactate shuttle mechanism, where glucose is taken up by astrocytes and converted to lactate, which is then oxidized by neurons^73^. In line with this model, the contribution of astrocytes to [^18^F]FDG signal has recently been elegantly demonstrated in the rodent brain after activation of astrocytic glutamate transport^74^. However, this model has been debated over the last decade after some animal evidence has shown that: i) glucose is taken up preferentially by neurons in awake behaving mice; ii) hexokinase, which catalyses the first enzymatic steps in glycolysis, is highly enriched in neurons as compared to astrocytes, in both mouse and human cortex^75^. Our brain cell-type enrichment analysis is consistent with this last model of glucose uptake by neurons by showing predominance of genes highly expressed in neurons. Interestingly, we also found enrichment for genes highly expressed in endothelial cells, which are known to uptake [^18^F]FDG using similar mechanisms to those used by neurons^76^. However, astrocytic genes were mostly anti-correlated with [^18^F]FDG (i.e. enrichment of astrocyte genes among genes negatively weighted). Whether this simply reflects species differences or the fact that our [^18^F]FDG scans were acquired at rest (while in the abovementioned rodent study^74^, [^18^F]FDG signal was measured in response to astrocytic glutamate transport) deserves further exploration. Altogether, these data support the plausibility of imaging transcriptomics in capturing patterns of constitutive gene expression that align well with different elements of the neuronal biology.

Our second set of analyses was focused on two tracers typically used as astrocyte probes and two MRI maps sensitive to myelin. Here, our findings broadly supported our hypothesis that by using imaging transcriptomics we would be able to identify patterns of gene expression consistent with astrocyte and oligodendrocyte genes, respectively. For both [^11^C]BU99008 and L-[^11^C]deprenyl-D2 we did find enrichment among the top genes positively associated with the regional distribution of these tracers for genes highly expressed in astrocytes and belonging to biological processes that are relevant to astrocytic biology, such as protein targeting (which is a key regulator of glycogen synthesis in astrocytes^77^) and RNA and ribonucleoproteins metabolism (which participate in local translation of transcripts in astrocytic peripheral processes^78^). In addition to astrocytes, we also found enrichment for genes highly expressed in some subclusters of neurons. This is not entirely surprising for the following reasons. [^11^C]BU99008 binds the imidazoline2 binding site (I2BS), which is thought to be expressed in glia and implicated in the regulation of glial fibrillary acidic protein^79,80^. Despite the lack of consensus regarding the nature of I_2_ receptors, cellular distribution studies revealed that I_2_BS are primarily located on the outer membrane of mitochondria and may be novel allosteric binding sites of monoamine oxidases (MAO) A and B^81-83^. Few studies have addressed the issue of which cell types in brain (neurones and/or glia) express I_2_BS. However, few different studies seem to agree that I_2_BS is expressed in both neurons and astroglia^84^. Similarly, L-[^11^C]deprenyl-D2 is a radiotracer which binds to MAO-B^85^. MAO-B is found in the brain primarily in non-neuronal cells such as astrocytes and radial glia, but immunohistochemistry studies have also reported the presence of MAO-B in neurons^86-88^.

For both MT ratio and myelin WC, we found enrichment for genes highly expressed in oligodendrocytes and OPCs, which matches the know sensitivity of these MRI markers to myelin^89,90^. However, we also found enrichment for other non-neuronal cells, such as microglia, astrocytes, pericytes and endothelial cells. This observation is intriguing and dovetails with recent controversies about the interpretation of MT and WC as specific markers for myelin, particularly in the grey matter^91-94^. For instance, the MT phenomena can occur in any large macromolecule with low molecular tumbling rate, making the measure sensitive to a variety of cellular processes (such as dendrites or myelin), but specific to virtually none^95^. Altogether, these data support the plausibility of imaging transcriptomics in capturing patterns of constitutive gene expression that align well with different elements of the astroglia and oligodendrocytes biology.

Our third set of analyses was conducted on four TSPO tracers and one tracer for Cox-1, which we hypothesized to be in line with the regional constitutive expression of genes involved in the neuroimmune response axis, such as those expressed in microglia or astrocytes. TSPO is expressed ubiquitously in the body and is used as a biomarker of neuroinflammation because its upregulation in inflammatory conditions is strongly localized to microglia and macrophages. Cyclooxygenase (COX) enzymes perform the rate-limiting step in the synthesis of inflammatory mediators such as prostaglandins and thromboxanes from arachidonic acid. The two main isoforms, COX-1 and COX-2 are present constitutively in the brain. COX-1 is predominantly localized in microglia in the brain. In contrast to our predictions, apart from [^11^C]PK11195, microglia and astrocyte genes were not among the top enrichment hits for all the other TSPO and Cox-1 tracers. Instead, for [^18^F]DPA174, the top enrichment hit was pericytes, but we also found enrichment for microglia genes; for [^18^C]PBR28 and [^18^C]ER176, the top enrichment hit was endothelial cells, but enrichment for microglia genes was not present. For [^11^C]PS13, the top enrichment hit was the excitatory neurons subcluster 3d, but enrichment for microglia was also present.

At first, these results might seem contradictory, but they need to be considered within the context of our recent increasing understanding of the biology of TSPO^96^ and methodological challenges that pervade in-vivo TSPO quantification in the human brain^97^. TSPO is an evolutionarily conserved protein localized in the outer mitochondrial membrane, which has been linked to various (but not mutually exclusive) physiological processes, such as cholesterol transport and steroid hormone synthesis, apoptosis and cell viability, redox processes and oxidative stress, and mitochondrial respiration and bioenergetics^96,98^. Recent evidence suggests that TSPO expression is spread across several brain cell-types and single cell quantifications of its constitutive expression suggest that basal TSPO mRNA expression is most abundant in ependymal cells, vascular endothelial cells, and microglia^99^; its expression is also detectable in neurons, even though in very low quantities and possibly modulated by neuronal activity^99^. Therefore, our findings for the TSPO tracers, namely a consistent enrichment for pericytes and endothelial cells, are broadly in line with this recent view of the cell-type relative expression distribution of TSPO. Only for [^11^C]PK11195, which is a first-generation tracer associated with low amounts of specific binding^97^, we identified microglia as the strongest enrichment hit; in contrast, for the other three second-generation tracers, which have demonstrated superior specificity ^97^, we did not. We also noted that while the TSPO gene was among the genes most positively correlated with the distribution of [^11^C]PK11195 (Z=3.08, Rank = 932/15,633), all the other three tracers were mostly unrelated to the distribution of TSPO mRNA ([^18^F]DPA174: Z=0.40, Rank=6866/15,633; [^18^C]PBR28: Z=-2.18, Rank=13,533/15,633; [^18^C]ER176: Z=-2.38, Rank=13,612/15,633). These findings are puzzling; however, admittedly, direct comparisons between tracers in this study are rather complex because of the use of different methods of quantification for the different tracers^97^, different imaging facilities and different subjects. In this work, we used templates where quantification was implemented as per the original respective publications, accounting for the different subject TSPO genetic polymorphisms by excluding low-affinity binders^100^. Here, we did not intend to identify which of these four tracers might perform better in capturing TSPO or examine the impact of the quantification approach in the correlations of different tracers with TSPO mRNA. We strongly believe such endeavour deserves its own comprehensive evaluation (for a thorough review about the complexity around TSPO quantification see Turkheimer et al.^101^). Instead, we examined different tracers to triangulate findings, which altogether dovetail with the known complexity of TSPO biology and quantification^96,97^. For Cox-1, the strongest enrichment hit was one subcluster of excitatory neurons, but as predicted we also found enrichment for microglia. This observation matches previous reports describing expression of Cox-1 in both neurons and microglia, but not astrocytes^102^. Moreover, we also note the enrichment for endothelial cells and pericytes, which aligns with descriptions of expression of Cox-1 in endothelial cells and its role in regulating the synthesis of prostacyclins in the cerebrovasculature^103,104^.

Our findings should be interpreted within the context of some limitations. First, given that most PET tracers do not have affinity for a single cell-type, our data should not be taken as an absolute validation of the imaging transcriptomics approach, since most likely tracer binding reflects contributions from more than one cell type. Instead, here, we examined whether the results of our enrichment analyses were plausible, given the known molecular nature of the tracers. Currently, we do not have access to dense phenotyping of the distribution of different brain cell types across the whole human brain, which admittedly would have been our first option for this work if existent. In the absence of this knowledge, PET provides only a practical approximation. Second, while still a general limitation of the field and not of this specific work, the AHBA whole-brain gene expression data derives only from six *post-mortem* adult brains (mean age = 43 y) and includes data in the right hemisphere from two donors, which led us to exclude the right hemisphere for the transcriptomic association analyses. Third, here we focused on comparisons between neuroimaging markers in the healthy adult human brain and constitutive gene expression in the healthy post-mortem brain. This does not detract that the strongest source of tracer binding might follow other biological pathways in pathology. A simple example is TSPO, which might be upregulated in microglia and astrocytes during neuroinflammation; in that case, if the disease process recruits the neuroimmune axis globally, then it is more likely that the TSPO distribution might better follow the constitutive regional distribution of astrocytes/microglia genes; while in the healthy brain, the signal might mirror better the endothelial component. In a similar way the radiotracer non displaceable non-specific binding might lead to spurious correlations. A simple example of this is represented by [^18^F]GE179, which has shown to suffer from low specific binding when used at baseline^105^. Fourth, by using constitutive gene expression in a small cohort of six post-mortem brains to infer associations with neuroimaging markers acquired in different cohorts, we are assuming that regional gene expression is a conserved canonical signature that generalizes well beyond the brain samples included in the AHBA. While we focused our analyses on probes that were selected to maximize differential stability across donors, six post-mortem brains are insufficient to make strong claims about the stability of gene expression across brains in humans. This might be a bigger concern for mobile brain cells that can dynamically move within the brain parenchyma, such as microglia, which participate in immunosurveillance even at rest^106^; this mobility might – at least theoretically – result in higher variability in gene expression across individuals and deserves further exploration. Fifth, no replicability analyses on independent datasets (same tracer, same experimental protocol and analysis, but different subjects) were performed. To our knowledge this work has been able to pull together an unprecedent number and varieties of brain scans. However, with the brain PET community recognising the importance of open data sharing (e.g. OpenNeuroPET, https://openneuro.org/pet)^107^, such endeavour might be feasible in the near future. Finally, our findings do show robustness to the specific choice of parcellation and set of single-cell genes used in our main analyses. For instance, we found large correlations between PLS1 genes weights between the analyses performed with the DK and AAL3 atlases; we also found that changing the set of genes used to define brain cell-types did not change our main conclusions considerably. However, it also true that both imaging and mRNA findings can be quite dependent on the particular methodology used to analyse the data. More work would in this topic should follow to ensure reproducibility and replicability.

In summary, our data supports the value and robustness of integrative imaging transcriptomics analyses in recovering plausible transcriptomic and cellular correlates of the regional distribution of a range of benchmark molecular imaging markers spanning different elements of the biology of the brain. The application of this indirect approach to bridge levels between gene expression, cells and macroscopical neuroimaging phenotypes holds the potential to improve our understanding of the biological pathways underlying regional variability in neuroimaging features. As a result, imaging transcriptomics could open new opportunities to expand the outputs of the use of neuroimaging tools in clinical research by: i) refining models of brain disease with the inclusion of more mechanistic biological information; ii) advancing the understanding of the role that genetic factors might play in brain regional vulnerability in brain disease; and iii) helping prioritizing targets for drug development or development of preclinical models with high translational value.

## List of Supplementary Materials

Supplementary methods: PET templates (full description).

Supplementary methods: Sensitivity analyses

Supplementary Figure S1. Correlations between neuroimaging markers.

Supplementary Table S1. Sensitivity analysis – brain parcellation.

Supplementary Figure S2. Sensitivity analysis – cortical + subcortical regions vs cortical regions only.

## Acknowledgments

We would like to thank all volunteers contributing data to this study.

## Author contributions

DM, MV, OD and FT designed the study; DM performed the data analysis and drafted the manuscript; AG wrote the python script provided with the manuscript; all authors discussed the findings, revised the manuscript for intellectual content and approved the final version of the manuscript.

## Appendix I PET templates references and contact points

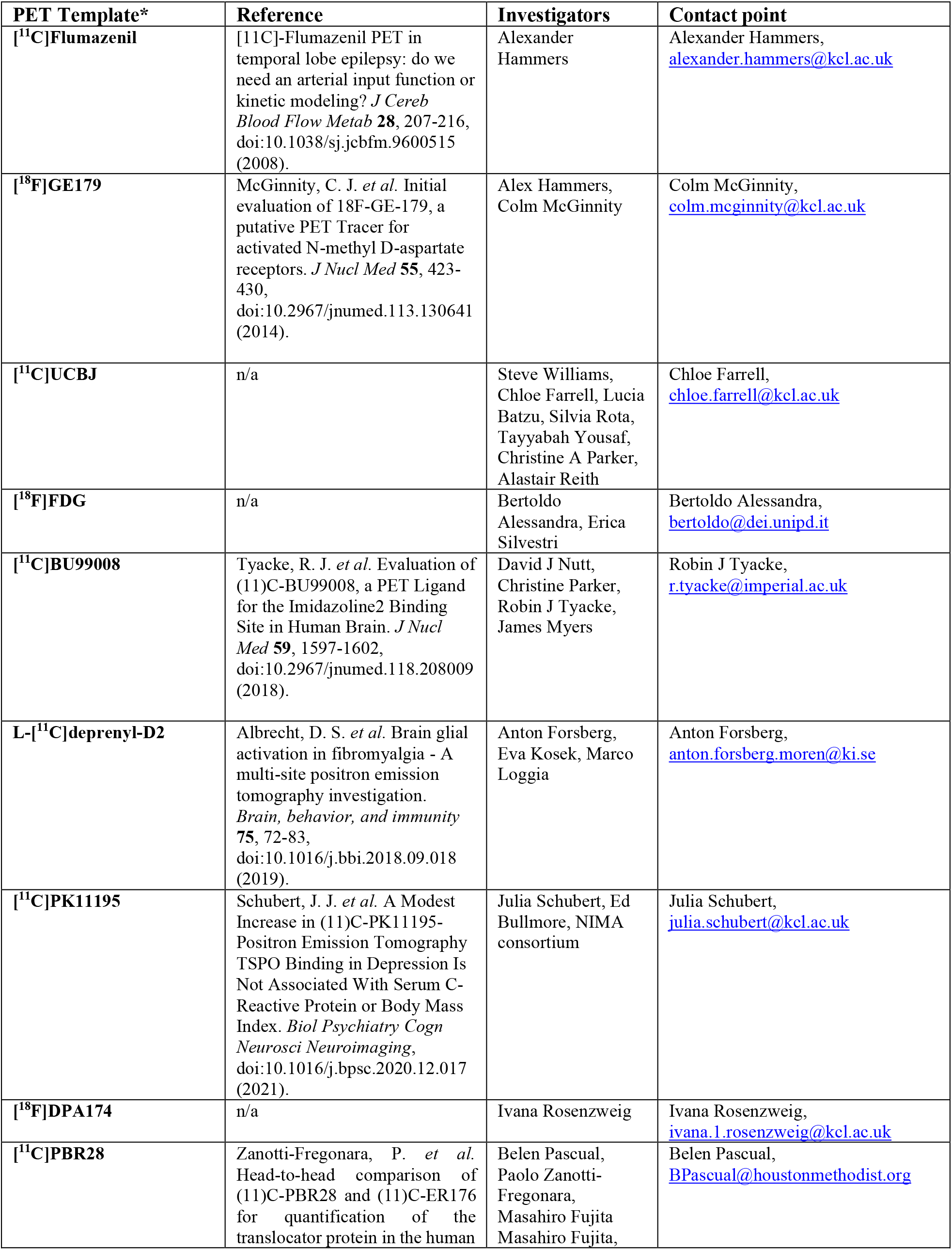
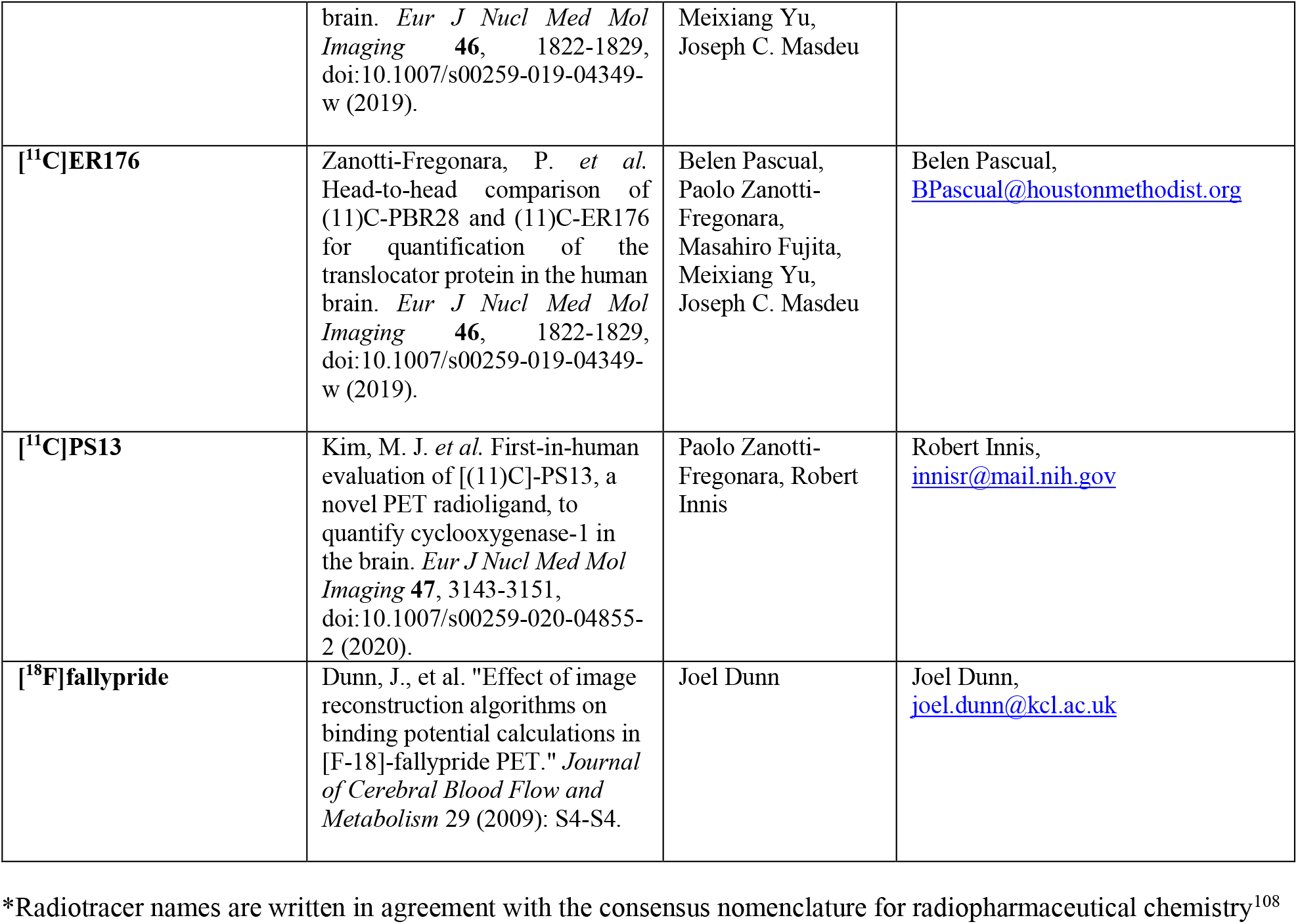

## Appendix II PET templates working group (in alphabetic order)

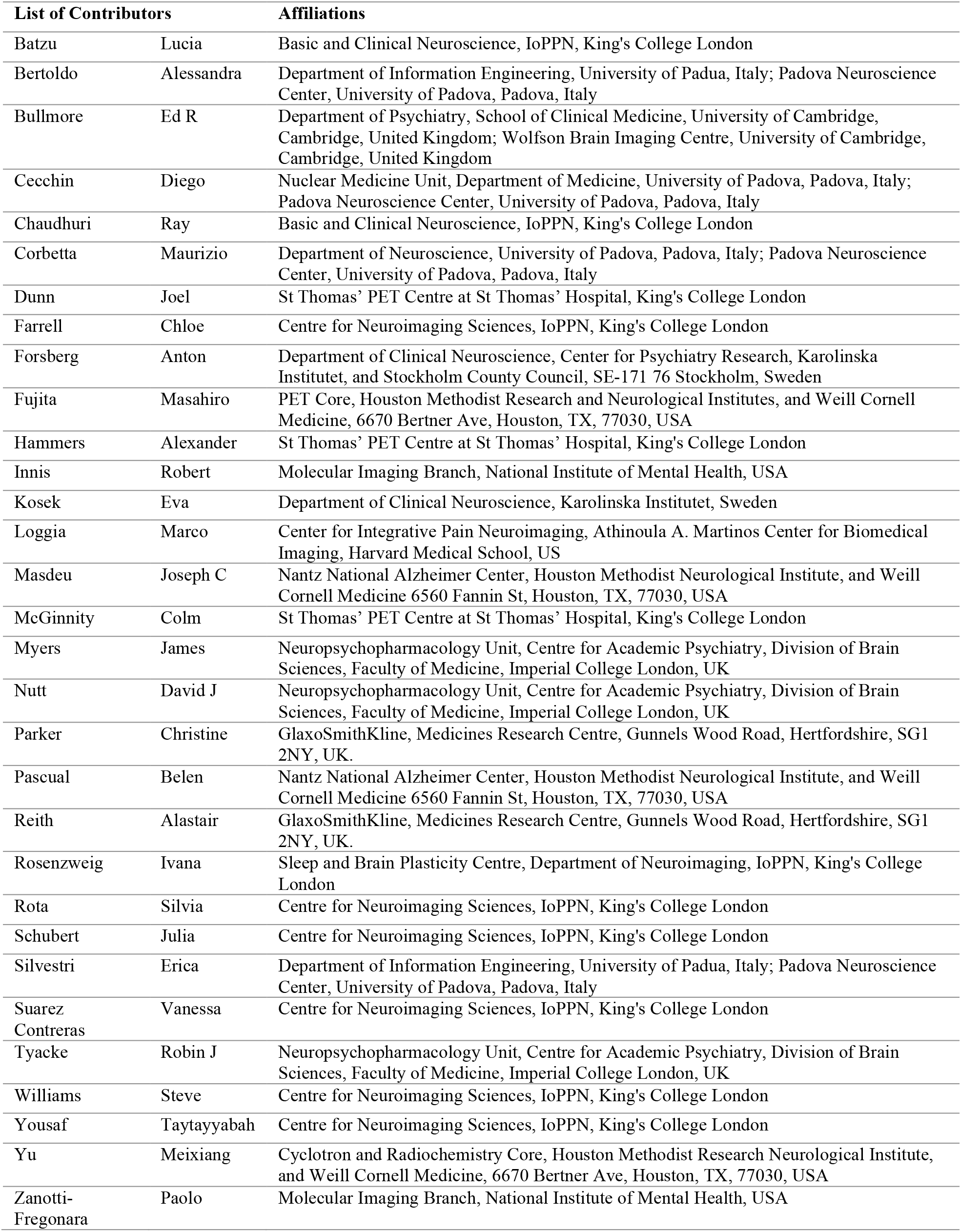

Appendix II – PET templates working group (in alphabetic order)
| List of Contributors |  | Affiliations |
| --- | --- | --- |
| Batzu | Lucia | Basic and Clinical Neuroscience, IoPPN, King's College London |
| Bertoldo | Alessandra | Department of Information Engineering, University of Padua, Italy; Padova Neuroscience Center, University of Padua, Padova, Italy |
| Bullmore | Ed R | Department of Psychiatry, School of Clinical Medicine, University of Cambridge, Cambridge, United Kingdom; Wolfson Brain Imaging Centre, University of Cambridge, Cambridge, United Kingdom |
| Cecchin | Diego | Nuclear Medicine Unit, Department of Medicine, University of Padua, Padova, Italy; Padova Neuroscience Center, University of Padua, Padova, Italy |
| Chaudhuri | Ray | Basic and Clinical Neuroscience, IoPPN, King's College London |
| Corbetta | Maurizio | Department of Neuroscience, University of Padua, Padova, Italy; Padova Neuroscience Center, University of Padua, Padova, Italy |
| Dunn | Joel | St Thomas' PET Centre at St Thomas' Hospital, King's College London |
| Farrell | Chloe | Centre for Neuroimaging Sciences, IoPPN, King's College London |
| Forsberg | Anton | Department of Clinical Neuroscience, Center for Psychiatry Research, Karolinska Institutet, and Stockholm County Council, SE-171 76 Stockholm, Sweden |
| Fujita | Masahiro | PET Core, Houston Methodist Research and Neurological Institutes, and Weill Cornell Medicine, 6670 Bertner Ave, Houston, TX, 77030, USA |
| Hammers | Alexander | St Thomas' PET Centre at St Thomas' Hospital, King's College London |
| Innis | Robert | Molecular Imaging Branch, National Institute of Mental Health, USA |
| Kosek | Eva | Department of Clinical Neuroscience, Karolinska Institutet, Sweden |
| Loggia | Marco | Center for Integrative Pain Neuroimaging, Athinoula A. Martinos Center for Biomedical Imaging, Harvard Medical School, US |
| Masdeu | Joseph C | Nantz National Alzheimer Center, Houston Methodist Neurological Institute, and Weill Cornell Medicine 6560 Fannin St, Houston, TX, 77030, USA |
| McGinnity | Colm | St Thomas' PET Centre at St Thomas' Hospital, King's College London |
| Myers | James | Neuropsychopharmacology Unit, Centre for Academic Psychiatry, Division of Brain Sciences, Faculty of Medicine, Imperial College London, UK |
| Nutt | David J | Neuropsychopharmacology Unit, Centre for Academic Psychiatry, Division of Brain Sciences, Faculty of Medicine, Imperial College London, UK |
| Parker | Christine | GlaxoSmithKline, Medicines Research Centre, Gunnels Wood Road, Hertfordshire, SG1 2NY, UK. |
| Pascual | Belen | Nantz National Alzheimer Center, Houston Methodist Neurological Institute, and Weill Cornell Medicine 6560 Fannin St, Houston, TX, 77030, USA |
| Reith | Alastair | GlaxoSmithKline, Medicines Research Centre, Gunnels Wood Road, Hertfordshire, SG1 2NY, UK. |
| Rosenzweig | Ivana | Sleep and Brain Plasticity Centre, Department of Neuroimaging, IoPPN, King's College London |
| Rota | Silvia | Centre for Neuroimaging Sciences, IoPPN, King's College London |
| Schubert | Julia | Centre for Neuroimaging Sciences, IoPPN, King's College London |
| Silvestri | Erica | Department of Information Engineering, University of Padua, Italy; Padova Neuroscience Center, University of Padua, Padova, Italy |
| Suarez Contreras | Vanessa | Centre for Neuroimaging Sciences, IoPPN, King's College London |
| Tyacke | Robin J | Neuropsychopharmacology Unit, Centre for Academic Psychiatry, Division of Brain Sciences, Faculty of Medicine, Imperial College London, UK |
| Williams | Steve | Centre for Neuroimaging Sciences, IoPPN, King's College London |
| Yousaf | Taytayyabah | Centre for Neuroimaging Sciences, IoPPN, King's College London |
| Yu | Meixiang | Cyclotron and Radiochemistry Core, Houston Methodist Research Neurological Institute, and Weill Cornell Medicine, 6670 Bertner Ave, Houston, TX, 77030, USA |
| Zanotti-Fregonara | Paolo | Molecular Imaging Branch, National Institute of Mental Health, USA |

