## supplementary material for "Imaging transcriptomics: Convergent cellular, transcriptomic, and molecular neuroimaging signatures in the healthy adult human brain"

**PET templates (full description)**

**[$^{11}$C]Flumazenil**: 13 healthy volunteers (2 females), ranging from 23 to 64 years (median age, 32 years) underwent a 90 min dynamic PET study with [$^{11}$C]Flumazenil using a 953b Siemens/CTI PET camera (CTI, Knoxville, Tenn., USA). Scans were acquired after injecting ~370 MBq of tracer intravenously, axially normalized, movement corrected and quantified as binding potential ($BP_{ND}$) maps using the simplified reference tissue model (SRTM) with input kinetics derived from the pons[1,2].

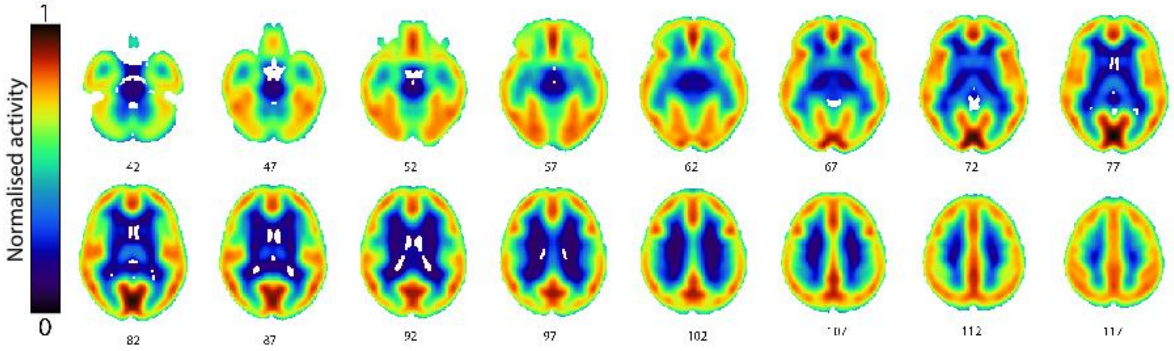


**[$^{18}$F]GE179**: 9 healthy volunteers (3 females), ranging from 25 to 62 years (median age, 37 years) underwent a 90 min dynamic PET scan using a ECAT EXACT3D 962 HR+ PET Camera (CTI/Siemens) after a median bolus injection of 186 MBq. Frames were reconstructed using a Fourier rebinning algorithm (FORE) and 2D filtered back projection, scans were then attenuation and motion corrected. Parametric $V_T$ maps were generated by voxelwise rank-shaping regularization of exponential spectral-analysis (RS-ESA)[3].

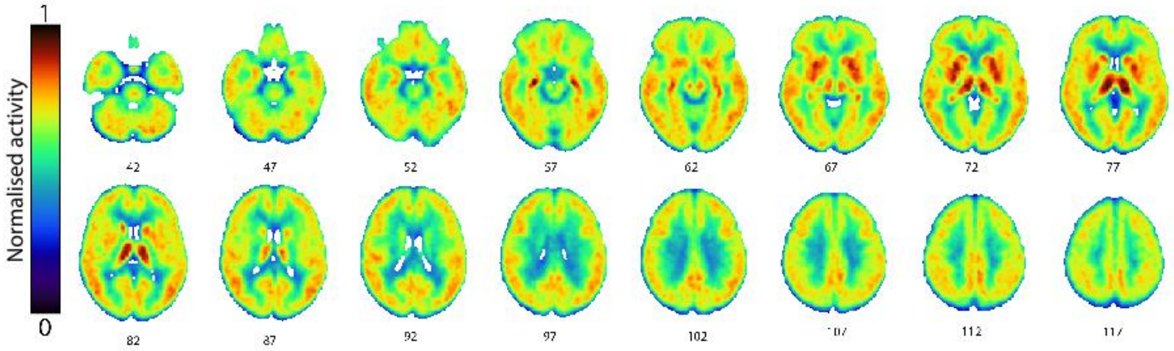

**$[^{11}C]$UCB-J**: 17 healthy volunteers (5 females, mean age ± SD, 60 ± 9 years) underwent a 90 min PET scan using a Siemens Biograph HiRez 6 PET/CT scanner after a bolus injection of 247 ± 47 MBq of $[^{11}C]$UCBJ. Scans were reconstructed by filtered back-projection, motion and attenuation corrected. Parametric $BP_{ND}$ maps were quantified by STRM using centrum semiovale as reference region[4,5].

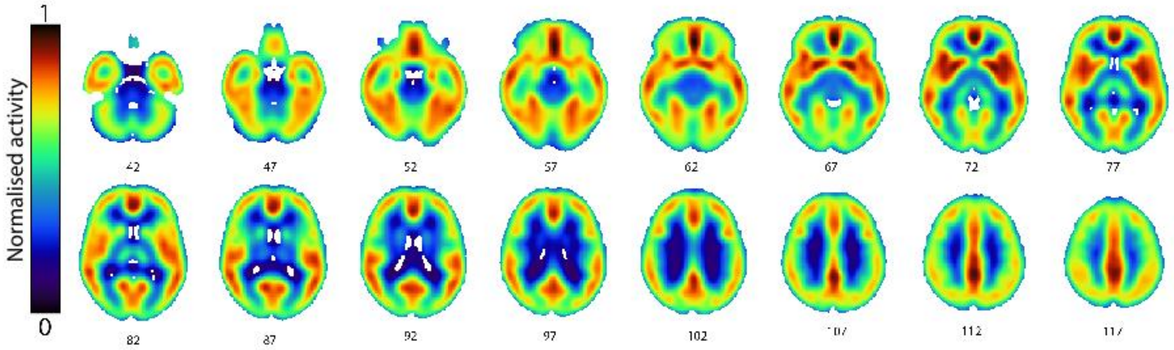


**$[^{18}F]$FDG**: 11 healthy volunteers (2 females, mean age ± SD, 52.9 ± 15 years) underwent a 20 min PET scan using a Siemens Biograph mMR PET/MR after a bolus injection of 209 ± 31.5 MBq of $[^{18}F]$FDG. Images were reconstructed by OP-OSEM, with 3 iterations, 21 subsets, attenuation corrected with UTE-derived maps and smoothed with a 4mm FWHM Gaussian filter. Final images were quantified with SUVR using whole brain white and grey matter as reference region.

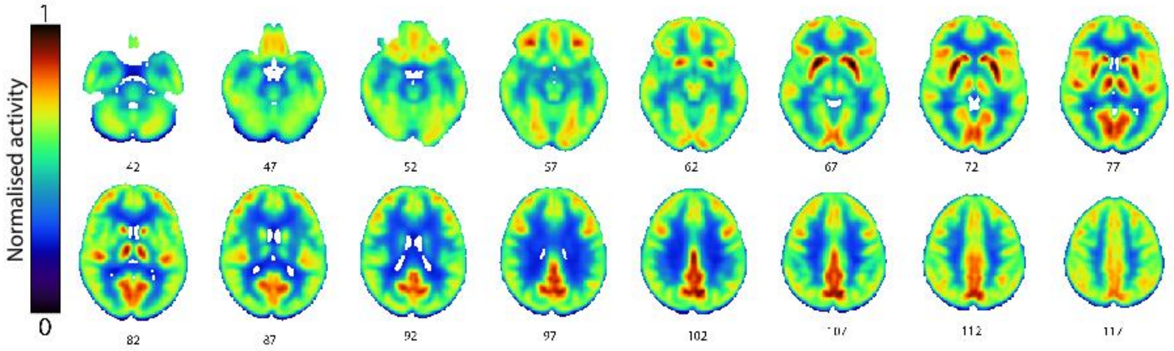

**[$^{11}$C]BU99008**: 13 healthy volunteers (1 female, mean age ± SD, 55 ± 7 years) underwent a 120 min dynamic PET scan using a Siemens HiRez Biograph 6 PET/CT scanner (Siemens Healthcare) after a bolus injection of 310 ± 13 MBq of [$^{11}$C]BU99008[6]. Images were reconstructed using filtered back projection, corrected for attenuation using CT based transmission and $V_T$ maps were quantified with Logan graphical analysis.

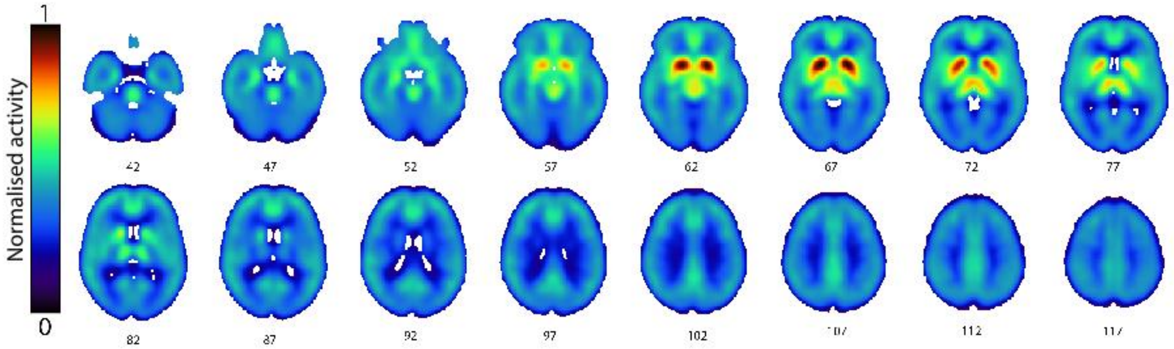


**L-[$^{11}$C]deprenyl-D2**: 11 healthy volunteers (11 females, mean age ± SD, 52± 8 years) underwent a 63 min dynamic PET scan using a Siemens High-Resolution Research Tomograph (HRRT, Siemens Molecular Imaging, Knoxville, TN, USA) after a bolus injection of 364 ± 46 MBq of L-[$^{11}$C]deprenyl-D2. Images were reconstructed by filtered back projection, with a 2mm Hanning filter, and were corrected for attenuation and scatter. Parametric maps were estimated by SUV, using the brain PET activity over the last 20 minutes of the acquisition [7].

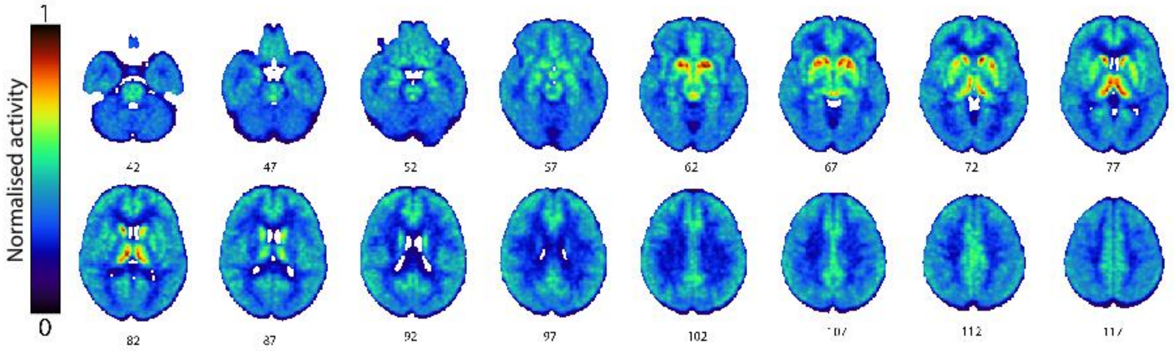

**[$^{11}$C]PK11195**: 25 healthy volunteers (14 females, mean age ± SD, 37 ± 8 years) underwent a 60 min dynamic PET scan using a GE SIGNA PET/MR (GE Healthcare, Waukesha, WI) after a bolus injection of 361 ± 53 MBq of [$^{11}$C]PK11195. Frames were reconstructed using time-of-flight (TOF) ordered subsets expectation maximization (OSEM), with 6 iterations, 16 subsets and no smoothing. Scans were motion corrected and $BP_{ND}$ maps were quantified via a simplified reference tissue model (SRTM) using a supervised clustering reference region approach[8].

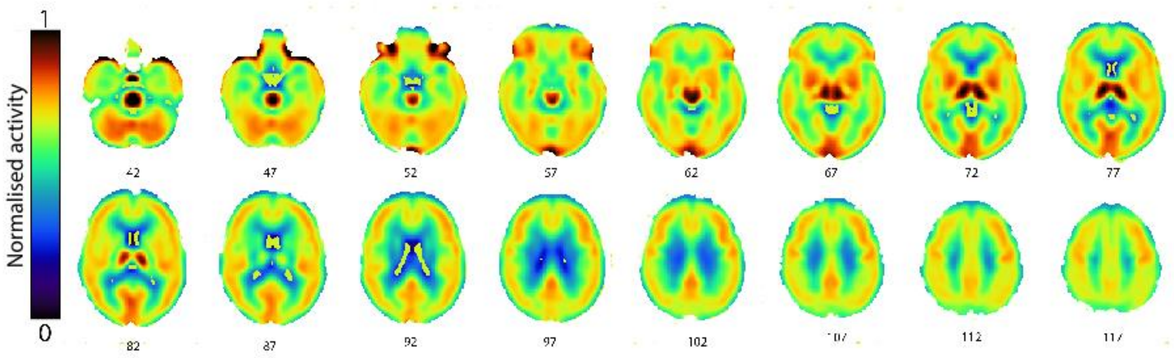


**[$^{18}$F]DPA174**: 10 healthy volunteers (all male, mean age ± SD, 34 ± 8 years) underwent a 60 min PET scan using a Siemens Biograph mMR PET/MR scanner after a bolus injection of 183 ± 20 MBq of [$^{18}$F]DPA174. Frames were reconstructed using time-of-flight ordered subset expectation maximization, with 6 iterations, 16 subsets and no smoothing. Images were motion corrected and quantified as binding potential ($BP_{ND}$) using a simplified reference tissue model and a supervised clustering reference region approach[9,10].

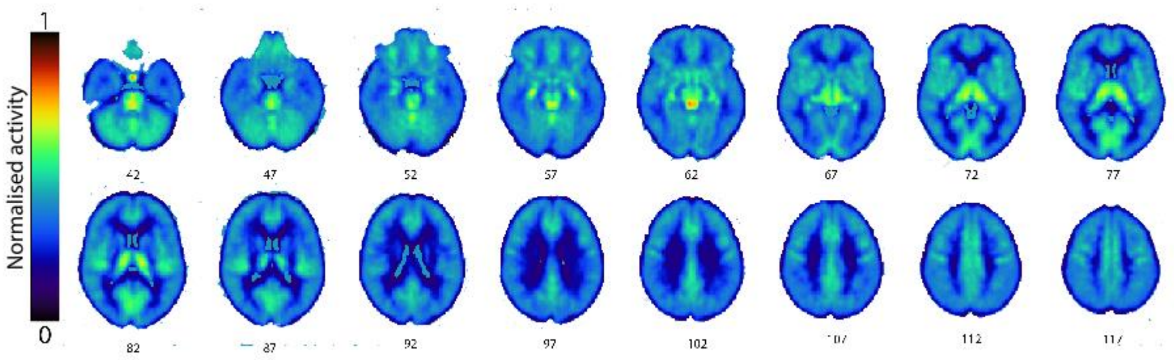

**[11C]PBR28**: 7 healthy volunteers (5 females, mean age ± SD, 68 ± 5 years) underwent a 90 min dynamic PET scan using a Philips Gemini TF 64 PET/CT scanner after a bolus injection of 743 ± 56 MBq of [$^{11}$C]PBR28. Voxelwise impulsive response function maps were quantified by Spectral Analysis Impulsive Response Function (SA-IRF)[11] computed at 90 minutes from tracer administration. Please note that the subjects were exactly the same participants used for the definition of [$^{11}$C]ER176 template. Individual TSPO tracer affinity was included as weight for the generation of population template. Only high- and medium-affinity binders were included.

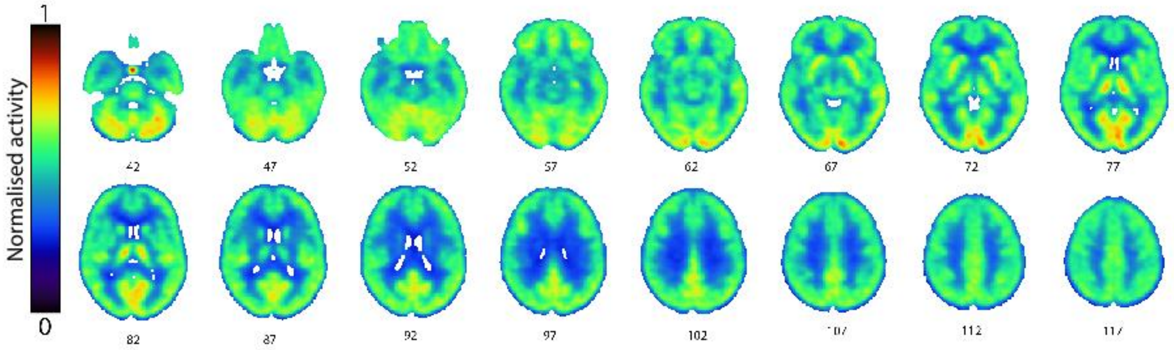


**[11C]ER176**: 7 healthy volunteers (the same for [$^{11}$C]PBR28 dataset) underwent a 90 min dynamic PET scan using a Philips Gemini TF 64 PET/CT scanner after a bolus injection of 723 ± 85 MBq of [$^{11}$C]ER176. Voxelwise impulsive response function maps were quantified by Spectral Analysis Impulsive Response Function (SA-IRF)[11] computed at 90 minutes from tracer administration. Individual TSPO tracer affinity was included as weight for the generation of population template.

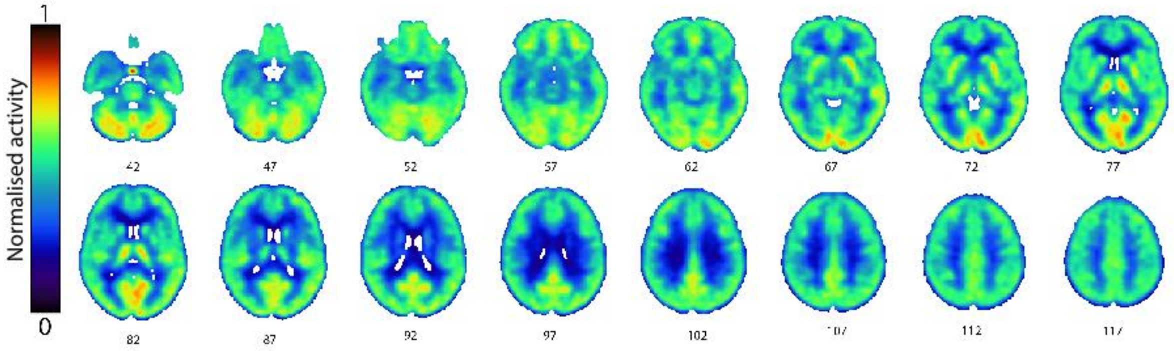

**[$^{11}$C]PS13**: 10 healthy volunteers (4 females, mean age ± SD, 29 ± 7 years) underwent a 120 min dynamic PET scan using a Siemens Biograph mCT (Siemens Healthineers, Erlangen, Germany) after a bolus injection of 690.2 ± 40.6 MBq of [$^{11}$C]PS13. Images were reconstructed with TOF OSEM with resolution recovery, with 3 iterations, 21 subsets and a 2mm FWHM Gaussian post-filter. Scans were motion corrected and $V_T$ maps were quantified by Logan graphical analysis[12].

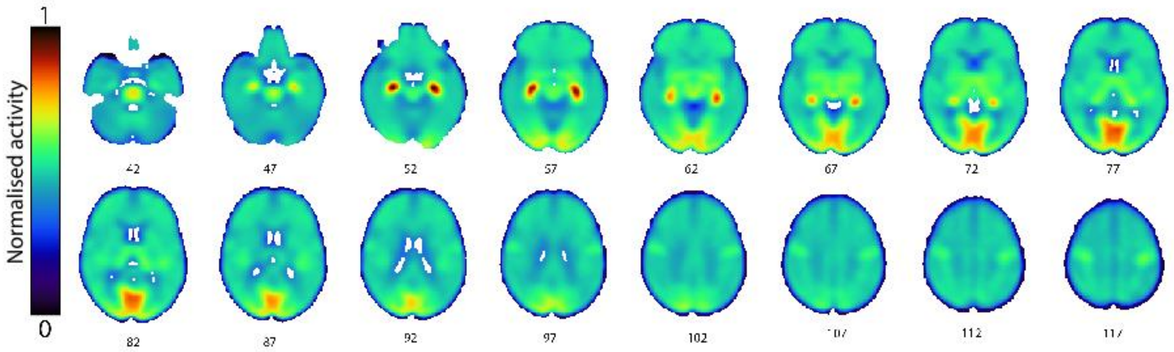


**MRI templates (full description)**

**Magnetization transfer**: 38 healthy volunteers (13 male; mean age = 34, range = 20–53). All MR imaging was performed on a Philips 3T Achieva scanner (Best, The Netherlands) using an eight-channel head RF array coil. The magnetization transfer imaging acquisition consisted of a 3D gradient echo sequence (TR = 85 ms, TE = 3.7 ms, 20 axial slices acquired at 5 mm slice thickness and reconstructed to 40 slices at 2.5 mm slice thickness, in-plane voxel size = 1 × 1 $mm^2$, FOV = 230 × 192 × 100 $mm^3$, $\alpha = 18°$, with and without an off-resonance RF pulse centered 1.1 kHz below the water frequency, sinc-gaussian envelope of duration = 15 ms, bandwidth = 190 Hz, amplitude = $2.3 \times 10^{-6}$ T).

**Myelin water content**: 31 healthy volunteers (16 females, mean age ± SD, 32.9 ± 6.8 years) underwent $T_1$ and $T_2^*$ and total water content MRI scans using a 3T Trio Siemens (Siemens Medical System, Erlangen, Germany) scanner. Full acquisition protocol and reconstruction is detailed in (Neeb et al 2008) and (Neeb 2012). Lesion corrected myelin water content data were smoothed via a 5x5 Gaussian kernel and linearly averaged[13].

**Sensitivity analyses:**

i) **Brain parcellation.** We examined how dependent our findings might be of the specific brain parcellation used (DK atlas), by repeating the PLS analysis but this time using both neuroimaging and transcriptomic data mapped to regions of the Anatomical Atlas Labelling, version 3. We used 64 regions of the left hemisphere, including 22 subcortical regions. As for the analyses with the DK atlas, we did not include cerebellum and brainstem. Then, for each marker, we correlated between analyses on different atlas the respective weights of all genes using Pearson correlations. Strong positive correlations in this case indicate that the determination of spatial alignment between gene and neuroimaging marker is minimally sensitive to the specific parcellation used in the main analyses.

ii) **Brain cell-types transcriptomic profile.** We examined how dependent our findings might have been of the specific transcriptomic profile of brain cell types used by repeating the same brain cell-type enrichment analyses but this time using another set of cell-types genes. For this analysis, we compiled data from four different single-cell studies (excluding *Lake et al.*) using *post-mortem* cortical samples in human postnatal subjects to avoid any bias based on acquisition methodology, analysis, or thresholding, as in a previous study[14]. To obtain gene sets for each cell type, categorical determinations were based on each individual study, as per the respective methods and analysis choices in the original paper. We generated a single omnibus gene list for each cell type by merging the study-specific gene lists, and then filtered it to retain only genes sampled in the AHBA. Two studies did not subset neurons into excitatory and inhibitory[15,16], and thus those gene sets were excluded from the cell-class assignment. All cell-type gene sets were available as part of the respective papers. The final pooled lists included sets of genes for astrocytes, microglia, endothelial cells, excitatory and inhibitory neurons, oligodendrocytes and OPCs.

iii) **Inclusion of cortical + subcortical regions vs cortical regions only**. A question often asked by researchers performing transcriptomic analyses is whether cortical and subcortical regions should be included in the same model, given the well-known strong anti-correlation between cortical and subcortical gene expression. This aspect has been previously examined by *Arnatkeviciute et al., 2019*[17], who suggested that cortical and subcortical data should be normalized separately to be considered simultaneously. Yet, studies tend to discard subcortical regions and focus on cortical regions only. When subcortical regions are of high interest, using cortical regions only might therefore be suboptimal. Therefore, we also investigated this assumption by performing imaging transcriptomics of [$^{18}$F]fallypride, a high affinity D2/D3 receptors antagonist, which binds highly in the basal ganglia but also return non negligible signal from the cortex [12]. Compared with other D2/D3 receptors antagonists radiotracers (e.g. [$^{11}$C]raclopride), [$^{18}$F] fallypride has higher affinity and higher signal-to-noise ratios in vivo and therefore provides more reliable quantitative measures of D2/D3 concentration including extra-striatal brain regions[13,14]. A [$^{18}$F]fallypride PET template was obtained by averaging six binding potential (BPND) whole brain maps acquired in healthy young volunteers (age range: 18-30 years)[15,16].We performed analyses considering cortical and subcortical regions together or cortical regions only and examined whether excluding subcortical regions would have a considerable impact on the ranking of the D2 receptor gene (*DRD2*) in our list of spatially associated genes.

**[$^{18}$F]fallypride PET template:** Subcortical activity

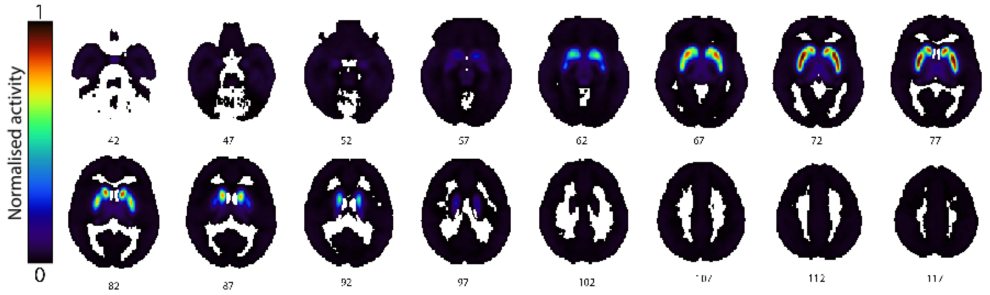


**[$^{18}$F]fallypride PET template***:* Cortical activity

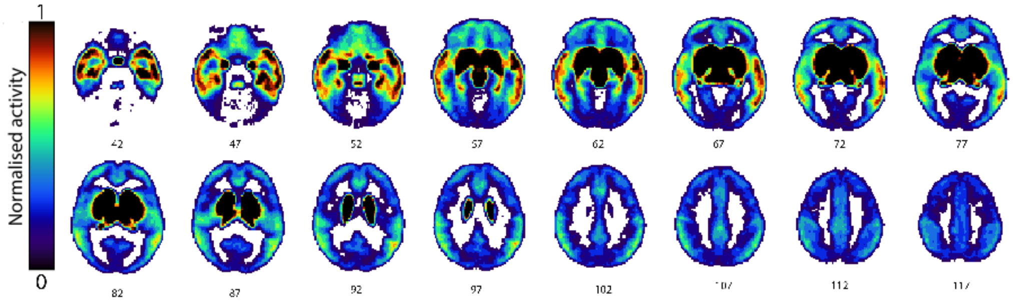

**Supplementary Figure S1. Correlations between neuroimaging markers.** Pairwise Pearson correlations between the Z-scores of the regional distribution of the neuroimaging markers. Significance was assessed with spin tests that account for the inherent spatial autocorrelation of the data. The symbol X indicates correlations that did not reach $p_{spin}$ <0.05. Abbreviations: MT – Magnetization transfer ratio; WC - Myelin water content.

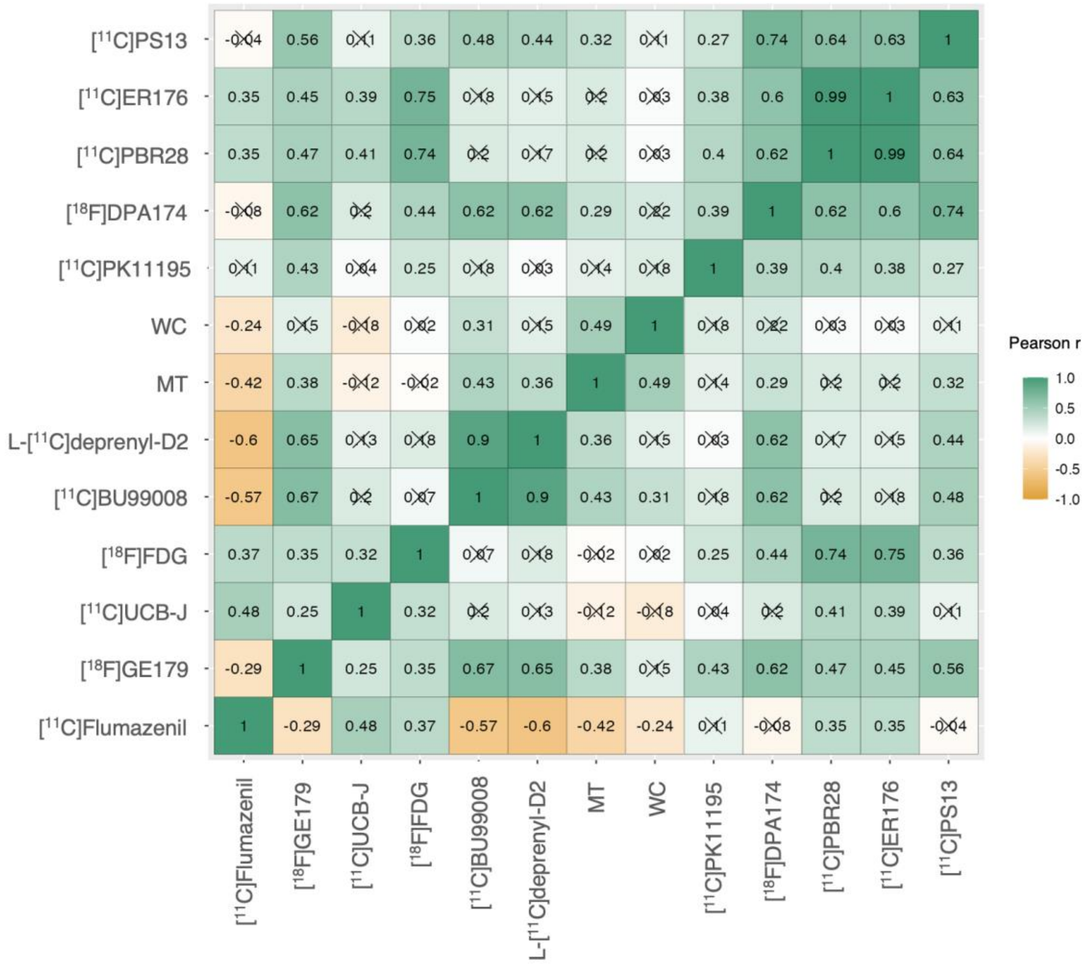

**Supplementary Table S1. Sensitivity analysis – brain parcellation.** Correlations between the weights of all genes as estimated in the analyses using the DK and AAL3 atlases. Strong positive correlations in this case indicate that the determination of spatial alignment between gene and neuroimaging marker is minimally sensitive to the specific parcellation used in the main analyses.

| **Marker** | **Pearson's correlation** | **CI$_{95\%}$** | **p** |
|---|---|---|---|
| [$^{11}$C]Flumazenil | 0.83 | 0.72; 0.94 | <0.001 |
| [$^{18}$F]GE179 | 0.79 | 0.64; 0.88 | <0.001 |
| [$^{11}$C]UCB-J | 0.91 | 0.85; 0.97 | <0.001 |
| [$^{18}$F]FDG | 0.9 | 0.86; 0.97 | <0.001 |
| [$^{11}$C]BU99008 | 0.94 | 0.89; 0.98 | <0.001 |
| L-[$^{11}$C]deprenyl-D2 | 0.94 | 0.88; 0.98 | <0.001 |
| MT | 0.82 | 0.72; 0.89 | <0.001 |
| WC | 0.83 | 0.72; 0.90 | <0.001 |
| [$^{11}$C]PK11195 | 0.85 | 0.76; 0.92 | <0.001 |
| [$^{18}$F]DPA174 | 0.89 | 0.80; 0.93 | <0.001 |
| [$^{18}$C]PBR28 | 0.92 | 0.87; 0.99 | <0.001 |
| [$^{18}$C]ER176 | 0.92 | 0.87; 0.99 | <0.001 |
| [$^{11}$C]PS13 | 0.88 | 0.79; 0.93 | <0.001 |

**Supplementary Figure S2. Sensitivity analysis – cortical + subcortical regions vs cortical regions only.** (A) Regional distribution of [$^{18}$F]fallypride; (B) Density plots showing the distribution of the weights of each gene, as determined in partial least square regression models based on cortical + subcortical regions (left) or cortical regions only (right). The black lines highlight the position of the *DRD2* gene in each distribution.

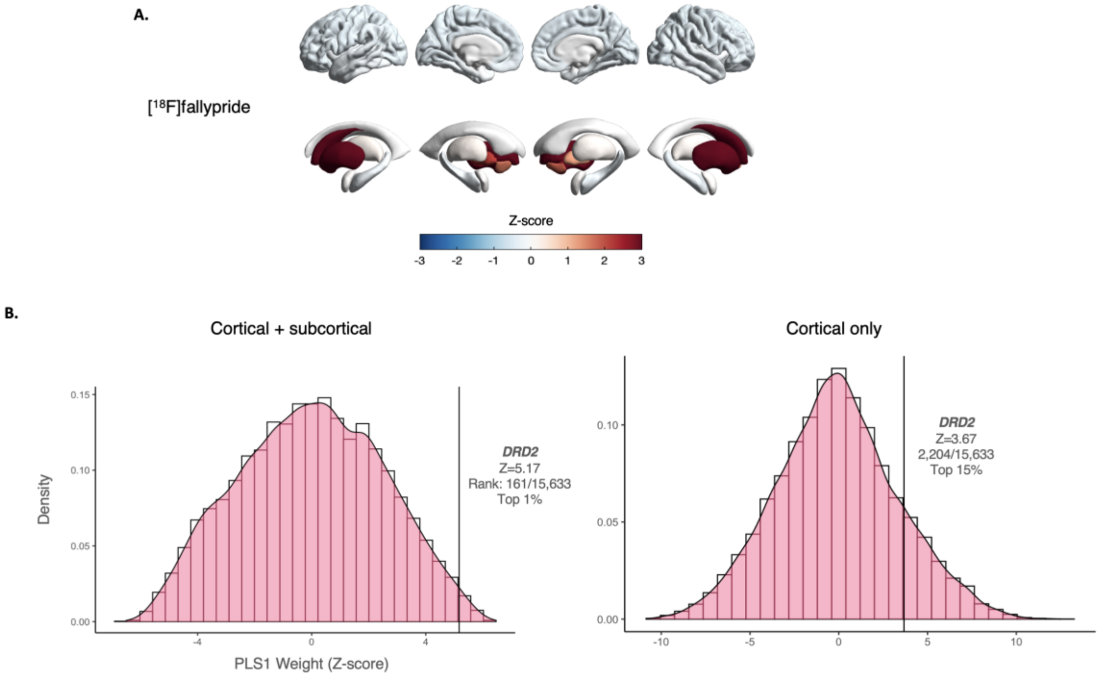
